# Large-scale brain network dynamics provide a measure of psychosis and anxiety in 22q11.2 deletion syndrome

**DOI:** 10.1101/551796

**Authors:** Daniela Zöller, Corrado Sandini, Fikret Işik Karahanoğlu, Maria Carmela Padula, Marie Schaer, Stephan Eliez, Dimitri Van De Ville

## Abstract

Prodromal positive psychotic symptoms and anxiety are two strong risk factors for schizophrenia in 22q11.2 deletion syndrome (22q11DS). The analysis of large-scale brain network dynamics during rest is promising to investigate aberrant brain function and identify potentially more reliable biomarkers. We retrieved and examined dynamics of large-scale functional brain networks using innovation-driven co-activation patterns (iCAPs) and probed into functional signatures of prodromal psychotic symptoms and anxiety. Patients with 22q11DS had shorter activation in cognitive brain networks and longer activation in emotion processing networks. Functional signatures of prodromal psychotic symptoms confirmed an implication of cingulo-prefrontal salience network activation duration and coupling. Functional signatures of anxiety uncovered an implication of amygdala activation and coupling, indicating differential roles of dorsal and ventral sub-divisions of anterior cingulate and medial prefrontal cortices. These results confirm that the dynamic nature of brain network activation contains essential function to develop clinically relevant imaging markers of psychosis vulnerability.

## Introduction

Schizophrenia is a strongly debilitating mental disorder both for affected individuals and in terms of societal cost (35, 31). Converging evidence suggests that schizophrenia is a progressive neurodevelopmental disorder, given that in most cases, sub-clinical psychiatric and cognitive symptoms of the disorder are present several years prior to the onset of a full-blown psychotic episode (35, 30, 45, 53, 73, 100). The neurodevelopmental model critically implies that earlier interventions might prove more effective in preventing the progression towards psychosis (53, 59). Hence extensive research has been devoted to characterizing the prodromal disease stage, also known as psychosis High-Risk State (30). In particular, clinical research has demonstrated that the presence of attenuated positive psychotic symptoms, operationalized in the Ultra-High-Risk criteria (99), confers a strongly increased 30-40% risk of developing psychosis (29). While current clinical management is based purely on clinical observation (63, 81), the identification of biomarkers of early psychosis could improve our understanding of the pathophysiology in its earliest disease stage (38). In this sense, the addition of imaging markers to the existing clinical diagnostic tools could allow the establishment of more precise biomarker-informed stages in the evolution of psychosis, which would give way to more targeted and effective therapeutic strategies and improved clinical outcomes (35, 59, 38).

Chromosome 22q11.2 deletion syndrome (22q11DS) is a neurodevelopmental disorder coming with a highly elevated risk for schizophrenia with a 30%-40% prevalence by adulthood (83). Most patients with 22q11DS are diagnosed already during childhood, which allows to characterize the earliest stages of schizophrenia’s disease course (35, 6). Similarly to the general population the presence of attenuated psychotic symptoms strongly increases the risk of psychosis in 22q11DS pointing to a common clinical trajectory with non-syndromic schizophrenia (82). Moreover, anxiety has emerged as another strong risk factor for psychosis in 22q11DS (32, 33). These clinical findings point to the particular importance of understanding the pathophysiology and characterizing biomarkers of attenuated psychotic symptoms and anxiety in 22q11DS.

Among the tools to characterize biomarkers, resting-state functional magnetic resonance imaging (rs-fMRI) has emerged as promising (78). FMRI provides the unique opportunity to non-invasively observe brain function, and the resting condition is especially well-suited in clinical populations because it requires minimal compliance from participants. Most studies on rs-fMRI in psychosis to date have used static functional connectivity (sFC); i.e., the correlation between the activation in different brain regions over the whole scanning time (91). However, a limitation of such static approaches is that they ignore the inherently dynamic nature of brain activity with potentially valuable information contained in dynamic changes of activation and connectivity over the scanning duration (11, 12, 71, 34, 43). In this perspective, dynamic approaches have the potential to identify more precise and more reliable biomarkers, and are particularly promising in schizophrenia, given the multiplicity of affected behavioral domains and brain circuits (10, 8, 24, 92, 91). Studies on dynamic brain function in schizophrenia point towards disrupted dynamic connectivity between several brain states, in particular of subcortico-cortical connectivity (15) and connections of the default mode network (DMN; 19, 60, 87, 75). The few studies to date investigating dynamic FC (dFC) in individuals at clinical high risk found reduced dFC of salience network (SN) and DMN (67) and stronger alterations in early schizophrenia patients than subjects at ultra high risk (18). These results underline the potential of dynamic brain function to improve our understanding of the pathophysiology in subjects at risk for schizophrenia.

Despite these promises of dynamic fMRI analysis, functional neuroimaging research in 22q11DS has so far mostly focused on static functional features (17, 80, 54, 84, 55), often targeting only specific functional networks such as the DMN (66, 85). The studies who explicitly investigated psychotic symptoms in 22q11DS showed correlations of DMN dysconnectivity with prodromal psychotic symptoms (17), as well as successful discrimination between patients at high vs. low risk based whole-brain rs-fMRI (80) and hypoconnectivity of DMN, SN, anterior cingulate cortex (ACC) and frontoparietal network (FPN; 84). Further, in the only two studies to date investigating a dynamic feature of brain function in 22q11DS – the variability of blood-oxygenation level dependent (BOLD) signals – we found widespread reductions in brain variability in 22q11DS (102), and reduced variability in the dorsal ACC in patients with higher prodromal psychotic symptoms (101). In general, aberrant function, but also structure of the ACC has been suggested as a neuroimaging marker for the development of psychosis in 22q11DS (65) and might reflect dysfunctional self-monitoring and salience processing, possible mechanisms for the emergence of psychosis (37).

There are multiple methods to investigate dynamic fMRI (71), many of which have already been applied in schizophrenia as outlined above (10, 18). In sliding-window dFC, changes in FC are extracted by computing FC in a temporal window that is shifted over time (11, 75). This method has also been extended to compute dFC between independent components obtained from independent component analysis (ICA). These approaches, however, are limited by the necessity to choose the window size, and are only able to detect relatively slow changes in FC (48). Alternatively, so-called first-order approaches rely on temporal clustering of fMRI frames to obtain patterns of co-activity, so-called “co-activation patterns” (CAPs) (50). Here, no minimum activation duration of a brain state needs to be specified, which allows to trace even fast changes is brain states. However, only one brain state (or CAP) can be active at a time point. To overcome these limitations, the recently introduced innovation-driven co-activation patterns (iCAPs) framework detects moments of significantly *changing* brain activity to extract large-scale brain networks and their dynamic properties (41, 43, 103). In other words, brain networks are retrieved from dynamic activation changes, rather than activation itself. In this way, the iCAPs framework allows to robustly retrieve spatially *and* temporally overlapping brain networks. Furthermore, it incorporates deconvolution of the hemodynamic response function as a prerequesite to network extraction, which makes it robust towards noise.

In this study, we complement the existing literature on dFC in schizophrenia by using iCAPs combined with multivariate pattern analysis to identify potential biomarkers for psychosis vulnerability in 22q11DS. We detect functional fingerprints of anxiety and positive prodromal symptoms, two symptoms that have emerged as reliable predictors of psychosis in 22q11DS (33, 82).

## Results

### Demographic characteristics

Table 1 shows demographic characteristics of the 78 patients with 22q11DS and 85 healthy controls (HCs) that were included in the current study. There were no significant differences in age, gender distribution and handedness of the two groups. IQ was significantly lower in patients with 22q11DS (p*<*0.001). Amongst patients, 55% met the criteria for one or multiple psychiatric diagnoses, while HCs were selected based on the absence of such diagnoses.

**Table 1:** Participants demographics. N/A = not applicable.

|  | HC | 22q11DS | p-value |
| --- | --- | --- | --- |
| Number of subjects (M/F) | 85 (36/49) | 78 (37/41) | 0.514 ($\chi^2$) |
| Age mean ± SD<br>(range) | 16.73 ± 5.85<br>(8.1-30.0) | 17.19 ± 5.37<br>(8.1-29.7) | 0.603 |
| Right handed* | 80.00% | 77.94% |  |
| IQ** | 110.12 ± 13.78 | 70.01 ± 12.41 | <0.001 |
| **N. subjects meeting criteria for**<br>**psychiatric diagnosis** | **N/A** | **43 (55%)** |  |
| Anxiety disorder | N/A | 9 |  |
| Attention deficit hyperactivity<br>disorder | N/A | 8 |  |
| Mood disorder | N/A | 5 |  |
| Schizophrenia or Schizoaffective Disorder | N/A | 4 |  |
| More than one psychiatric disorder | N/A | 17 |  |
| **N. subjects medicated** |  |  |  |
| Methylphenidate | 0 | 9 |  |
| Anitpsychotics | 0 | 3 |  |
| Anticonvulsants | 0 | 1 |  |
| Antidepressants | 0 | 1 |  |
| More than one class of medication | 0 | 3 |  |

\* Handedness was measured using the Edinburgh laterality quotient, right handedness was defined by a score of more than 50. \*\* IQ was measured using the Wechsler Intelligence Scale for Children-III (95) for children and the Wechsler Adult Intelligence Scale-III (96) for adults.

### Extracted spatial maps correspond to known resting-state networks

We applied the iCAPs framework to rs-fMRI scans of all subjects, including both HCs and patients with 22q11DS. We identified 17 iCAPs corresponding to well-known resting-state networks (see Figure 1 and Supplementary Table S2). See Materials and Methods for details on iCAPs analysis and the definiting of the optimum number of networks based on consensus clustering (62) (see also Supplementary Figures S1 and S2). The obtained networks included sensory-related networks such as primary visual (PrimVIS1 and PrimVIS2; iCAP1 and iCAP8), secondary visual (SecVIS, iCAP2), auditory/sensorimotor (AUD/SM, iCAP13) and sensorimotor (SM, iCAP6) networks. The default mode network (DMN) was decomposed into anterior (aDMN, iCAP10), posterior (pDMN, iCAP11) and precuneus/ventral DMN (PREC/vDMN, iCAP7). There were two attention-related iCAPs; i.e., fronto-parietal network (FPN, iCAP9) and visuospatial network (VSN, iCAP12). Two iCAPs included regions commonly considered as the salience network: the anterior insula (aIN, iCAP3) and dorsal anterior cingulate cortex together with dorso-lateral prefrontal cortex (dACC/dlPFC, iCAP5). The remaining iCAPs comprised a language network (LAN, iCAP4), inferior temporal and fusiform (iTEMP/FUS, iCAP14), amygdala and hippocampus (AMY/HIP, iCAP15), orbitofrontal cortex (OFC, iCAP16) and prefrontal cortex (PFC, iCAP17).

**Figure 1:**
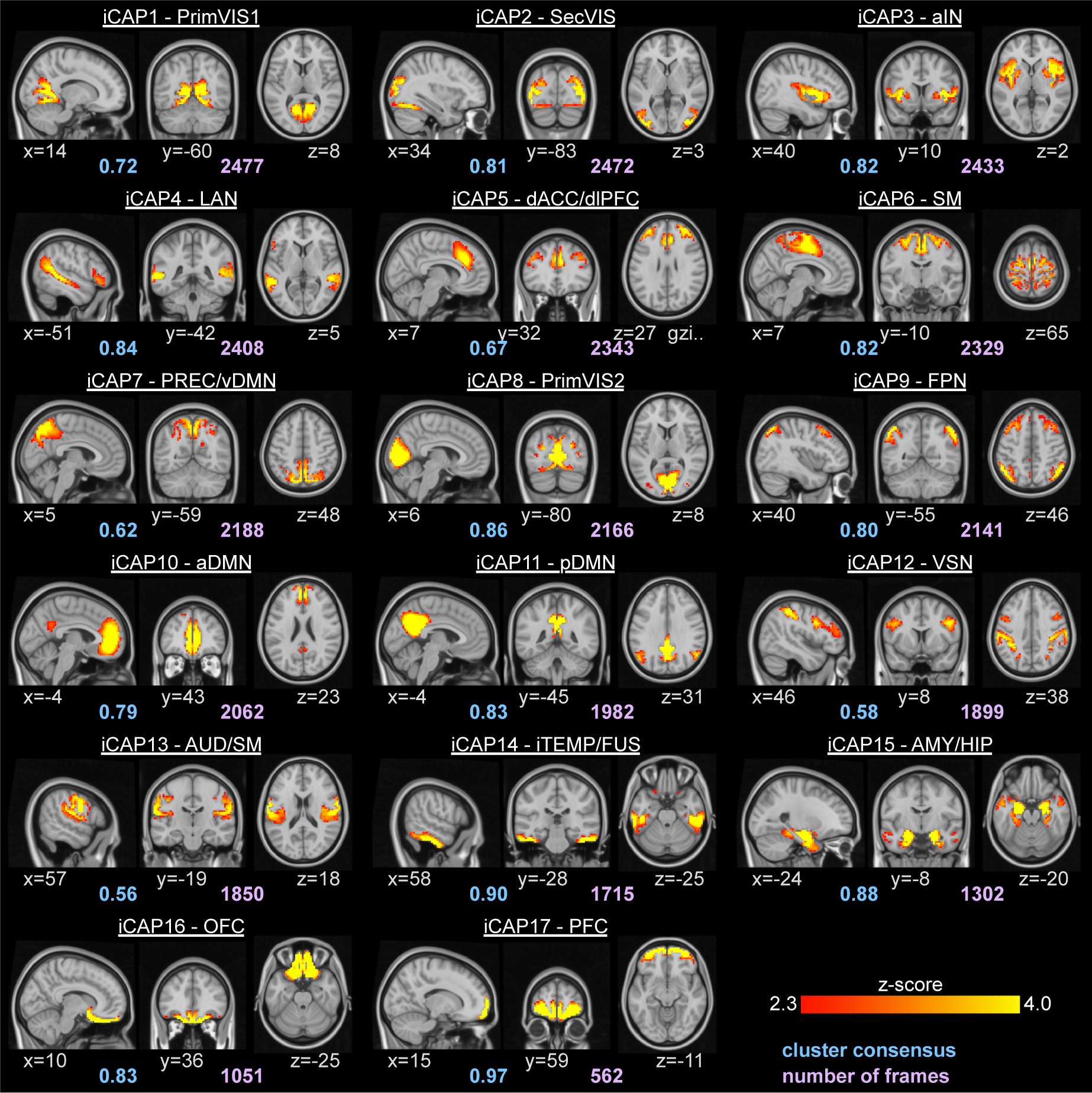
Spatial patterns of the 17 iCAPs retrieved from all subjects, including both HCs and patients with 22q11DS. The locations denote displayed slices in MNI coordinates. Blue values denote the average consensus of each cluster, purple values indicate the total number of innovation frames that were assigned to this cluster. PrimVIS1 – primary visual 1, SecVIS – secondary visual, aIN – anterior insula, LAN – language network, dACC/dlPFC – dorsal anterior cingulate cortex/dorsolateral prefrontal cortex, SM – sensorimotor, PREC/vDMN – precuneus/ventral default mode network, PrimVIS2 – primary visual 2, FPN – fronto-parietal network, aDMN – anterior default mode network, pDMN – posterior default mode network, VSN – visuospatial network, AUD/SM – auditory/sensorimotor, iTEMP/FUS – inferior temporal/fusiform, AMY/HIP – amygdala/hippocampus, OFC – orbitofrontal cortex, PFC – prefrontal cortex.

### Patients with 22q11DS show altered activation and coupling of iCAPs

Our first objective was to probe into alterations of the identified networks’ temporal properties in patients with 22q11DS. First, we investigated in which networks patients with 22q11DS had aberrant activation duration. Then, given the neurodevelopmental nature of 22q11DS, we tested whether these alterations might be related to aberrant development with age. Finally, we looked further into aberrant coupling duration between networks, which was assessed as positive coupling duration (same-signed co-activation between two iCAPs) or anti-coupling duration (differently-signed co-activation between two iCAPs).

### Altered duration of iCAPs’ activation

Figure 2 shows duration for all 17 iCAPs in percentage of total non-motion scanning time. Median total activation time ranged from 34.36 % for LAN (4) to 1.54 % for PFC (17). Patients with 22q11DS had significantly shorter activation of dACC/dlPFC (5), PrimVIS2 (8), FPN (9), aDMN (10) and pDMN (11) and significantly longer activation of SM (6), iTEMP/FUS (14), AMY/HIP (15) and OFC (16).

**Figure 2:**
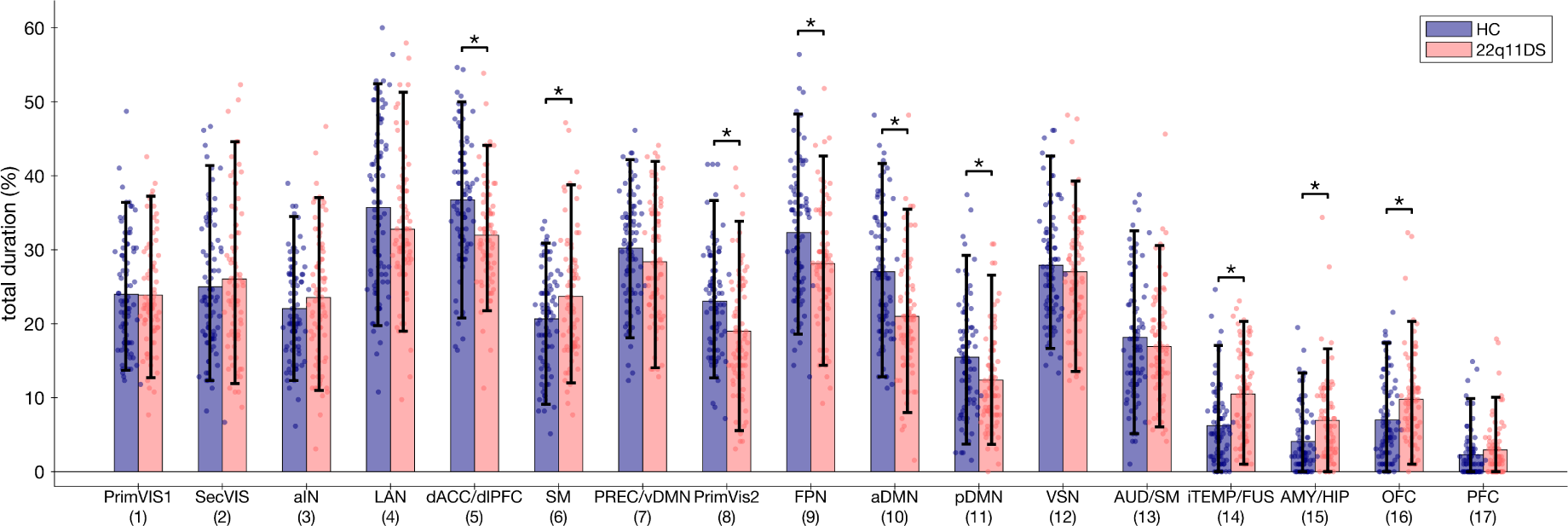
Statistics of total temporal duration for each iCAP. P-values are FDR-corrected for the 17 multiple comparisons and age, gender and motion were included as covariates. Significant group differences (p*<*0.05) were marked with an asterisk. Error bars indicate bootstrapping 5th to 95th percentiles. Single-subject duration measures were included as scatterplots. Corresponding test statistics (p-values, effect size) can be found in Supplementary Table S3.

### Relationship between age and duration of iCAPs’ activation

To assess the association between age and duration of iCAPs’ activation we conducted Partial Least Squares Correlation (PLSC; 46) – a technique that provides multivariate association between variables. PLSC is based on the extraction of principal components of correlation between a set of behavioral measures and a set of brain activation measures. It results in correlation components that are associated with brain and behavior weights indicating the contribution of each respective measure to the multivariate correlation (see Materials and Methods for more details).

We conducted PLSC with age as behavioral variable and duration of the nine altered iCAPs (figure 2) as brain variables. There was one significant correlation component (p=0.04) showing an opposite age-effect in HCs and patients with 22q11DS (age-weights: *−*0.44*±*0.2 in HCs, 0.89*±*0.21 in 22q11DS, see Supplementary Figure S3). In three networks (dACC/dlPFC – 5, PrimVIS2 – 8, FPN – 9), we found declining duration with age in patients with 22q11DS, but increasing duration with age in HCs. These three iCAPs had also globally shorter activation in 22q11DS. Contrarily, two networks (iTEMP/FUS – 14, AMY/HIP – 15) had increasing duration with age in 22q11DS and declining duration with age in HCs. Both of these networks had also globally longer activation in 22q11DS. Together, these results suggest that the alterations in these five networks are emerging over age in patients with 22q11DS.

### Alterations in temporal coupling of networks

To go beyond pure investigation of activation, we next probed into alterations of brain network interactions. Specifically, we analyzed the duration of positive coupling (co-activation with same sign) or anti-coupling (co-activation with opposite sign) between between all pair-wise combinations of iCAPs. Figure 3 shows significant group differences in iCAPs’ coupling. For several networks the duration of coupling was longer in patients with 22q11DS than in controls. This was true for 6 positive couplings and 13 anti-couplings. Fewer networks had shorter duration of coupling in patients with 22q11DS (1 positive coupling, 5 anti-couplings). Globally, alterations were more numerous for anti-couplings (25 in total) than for positive couplings (6 in total).

### Functional signature of positive psychotic symptoms

In order to investigate the behavioral relevance of this aberrant activation and coupling in 22q11DS, we conducted behavior PLSC including positive symptoms items deducted from the Structured Interview of Prodromal Symptoms (SIPS; 61). We first investigated iCAPs’ activation duration, followed by a second PLSC further testing for the implications of positive couplings and anti-couplings of the dACC/dlPFC (5) network.

### Altered iCAPs’ duration is associated with psychotic symptoms in patients with 22q11DS

PLSC analysis including positive SIPS items and iCAPs’ activation duration of the nine altered iCAPs (see figure 2) resulted in one significant correlation component (p=0.05, see figure 4A). The duration of dACC/dlPFC (5), FPN (9) and iTEMP/FUS (14) was positively correlated with all five positive psychotic symptoms.

### Altered couplings of dACC/dlPFC are associated with psychotic symptoms in patients with 22q11DS

Next, we investigated the relevance of couplings for psychotic symptoms. For this, we selected the dACC/dlPFC (5) network based on its appearance in the previous analysis (see previous paragraph “Altered iCAPs’ duration is associated with psychotic symptoms in patients with 22q11DS”), as well as literature associating ACC alterations with psychosis in 22q11DS (65). We included coupling time of dACC/dlPFC (5) with iCAPs that had altered couplings (iCAPs 3, 13, 14 and 15; see figure 3) and with iCAPs whose duration was significantly correlated with psychotic symptoms (iCAPs 9 and 14; see figure 4A).

**Figure 3:**
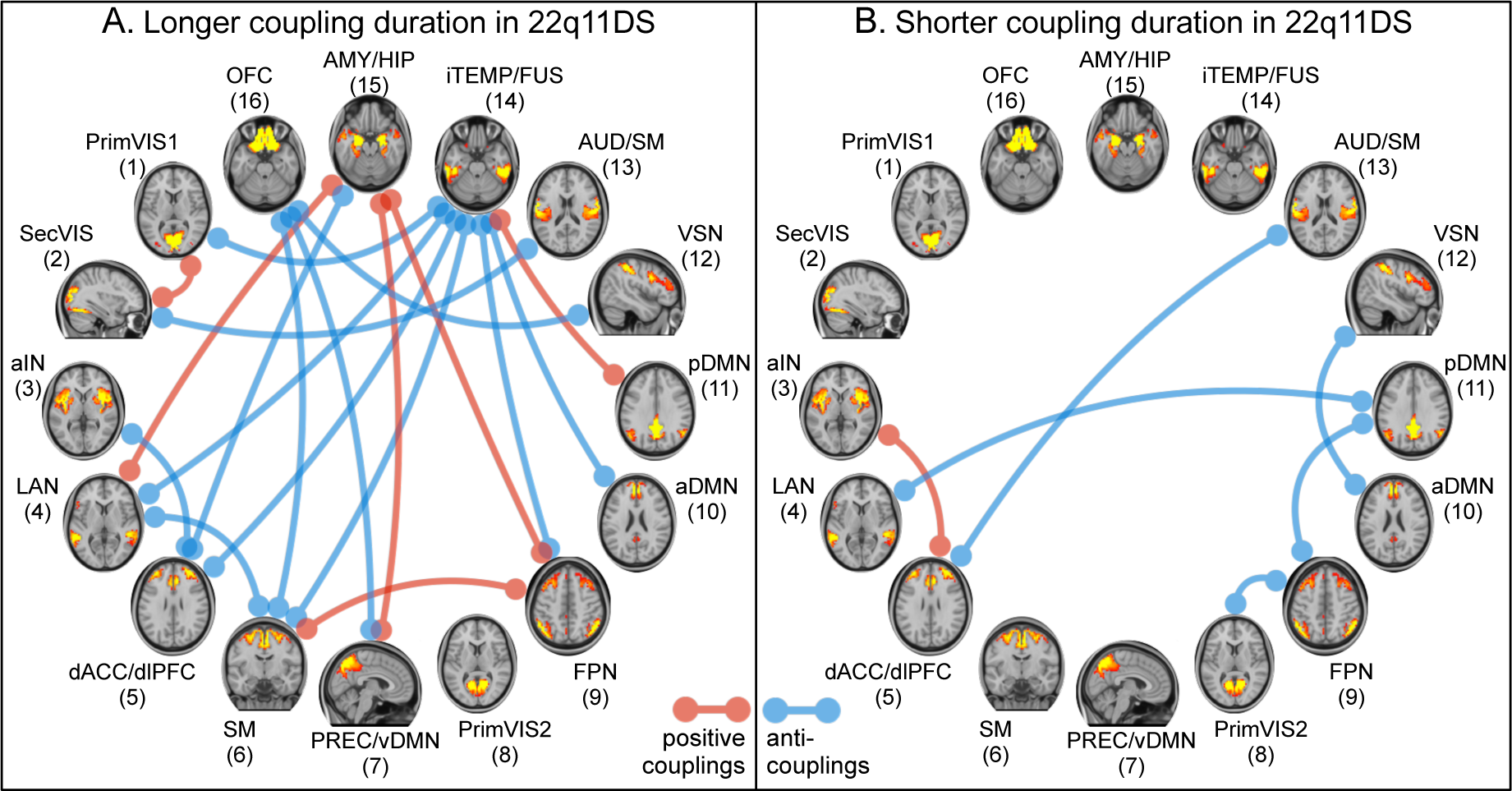
Significant duration differences of positive couplings (red) and anti-couplings (blue) between patients with 22q11DS and HCs. A) Couplings with significantly longer duration in 22q11DS. B) Couplings with significantly shorter duration in 22q11DS. Couplings were measured in terms of percentage of total scanning time, or in percentage of the joint activation time of the two respective iCAPs (Jaccard score). We here show only differences that were significant in both coupling measures. Underlying group comparison statistics can be found in Supplementary Figure S4 and Supplementary Table S4.

**Figure 4:**
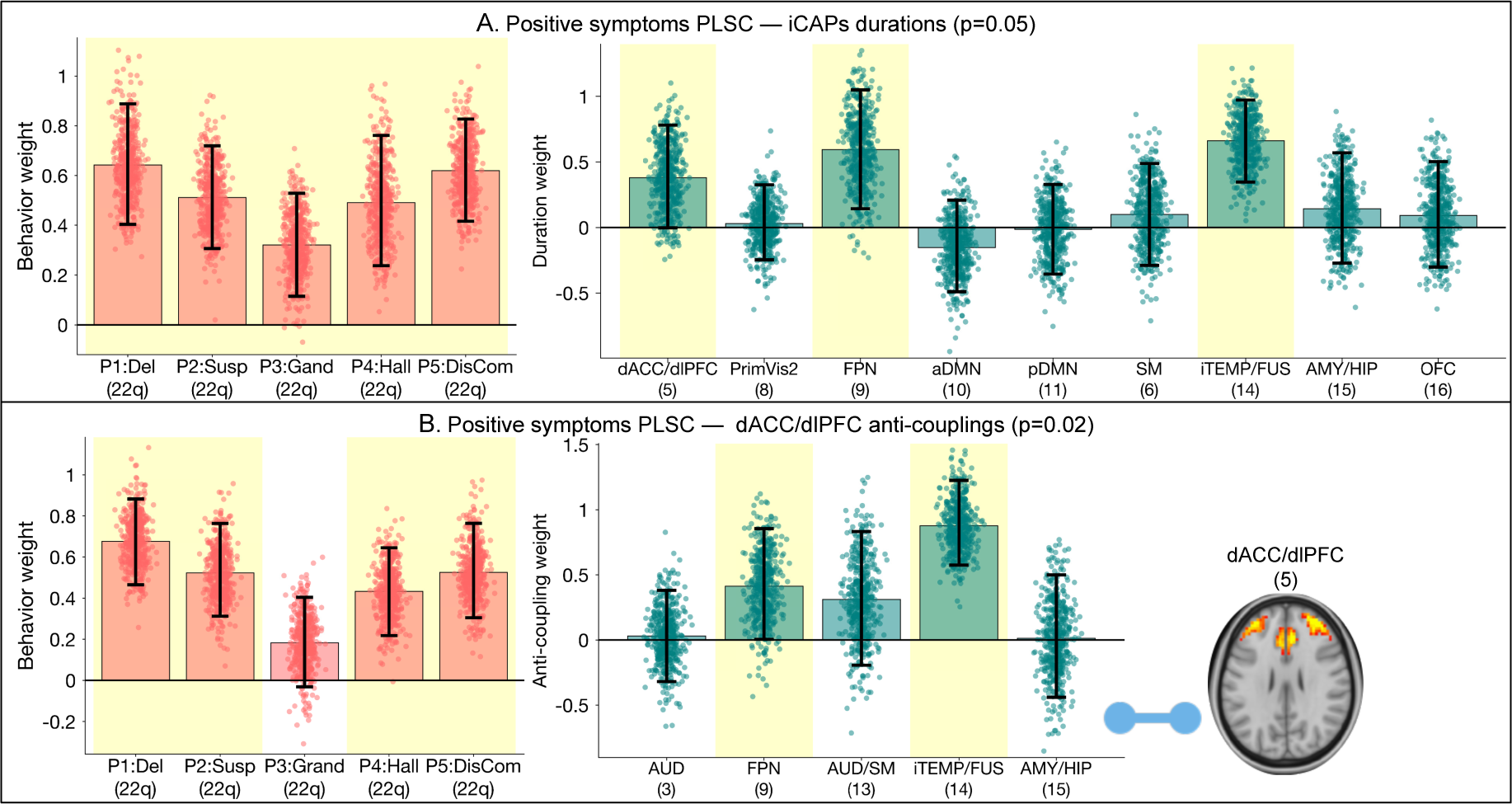
PLSC results for positive psychotic symptoms (Five SIPS items:Delusions, Suspiciousness, Grandiosity, Hallucinations and Disorganized Communication) in patients with 22q11DS. A) Behavior weights and brain weights for PLSC including duration of nine iCAPs with altered duration in 22q11DS. There is a positive correlation of positive psychotic symptoms with duration of dACC/dlPFC (5), FPN (9) and iTEMP/FUS (14). B) Behavior weights and brain weights for PLSC including anti-couplings of dACC/dlPFC (5) that were altered in 22q11DS. Longer anti-coupling of dACC/dlPFC (5) with FPN (9) and iTEMP/FUS (14) is associated with higher positive symptoms. Error bars indicate bootstrapping 5th to 95th percentiles, robust results were indicated by yellow background. Exact values of bootstrap mean and 5-95 percentiles are reported in Supplementary Table S5. PLSC results for positive couplings were not significant (p=0.6) and are thus not reported here.

We first conducted a PLSC for anti-coupling time between dACC/dlPFC (5) and these networks and found one significant correlation component (p=0.02, see figure 4B) showing an association between higher positive symptoms and longer anti-coupling of dACC/dlPFC (5) with FPN (9) and iTEMP/FUS (14). Thus, not only the activation of dACC/dlPFC (5) by itself is associated with higher symptoms severity, but also its interaction with other networks, namely its anti-coupling time with FPN (9) and iTEMP/FUS (14).

A second PLSC analysis for positive coupling time between dACC/dlPFC (5) and these networks did not give any significant correlation component (p=0.58).

### Functional signature of anxiety

Finally, we conducted similar analyses to investigate dynamic brain network alterations associated with anxiety, another behavioral risk factor for psychosis in 22q11DS besides prodromal symptoms. Anxiety scores were obtained from the Child Behavioral Checklist (CBCL) (1) in children, and the Adult Behavioral Checklist (ABCL) in adults (see also Materials and Methods). Again, we first probed into iCAPs’ activation duration, followed by a further analysis of positive couplings and anti-couplings – this time of the AMY/HIP (15) network.

### Altered iCAPs’ duration is associated with anxiety in patients with 22q11DS and HCs

We performed PLSC analysis between CBCL/ABCL anxiety scores in 22q11DS and HCs and iCAPs’ duration, again including the nine iCAPs with altered duration (see figure 2). There was one significant correlation component (p=0.03, see figure 5A). Both in HCs and patients with 22q11DS, longer activation of PrimVIS2 (8), iTEMP/FUS (14) and AMY/HIP (15) and shorter activation of aDMN (10) were associated with higher anxiety.

**Figure 5:**
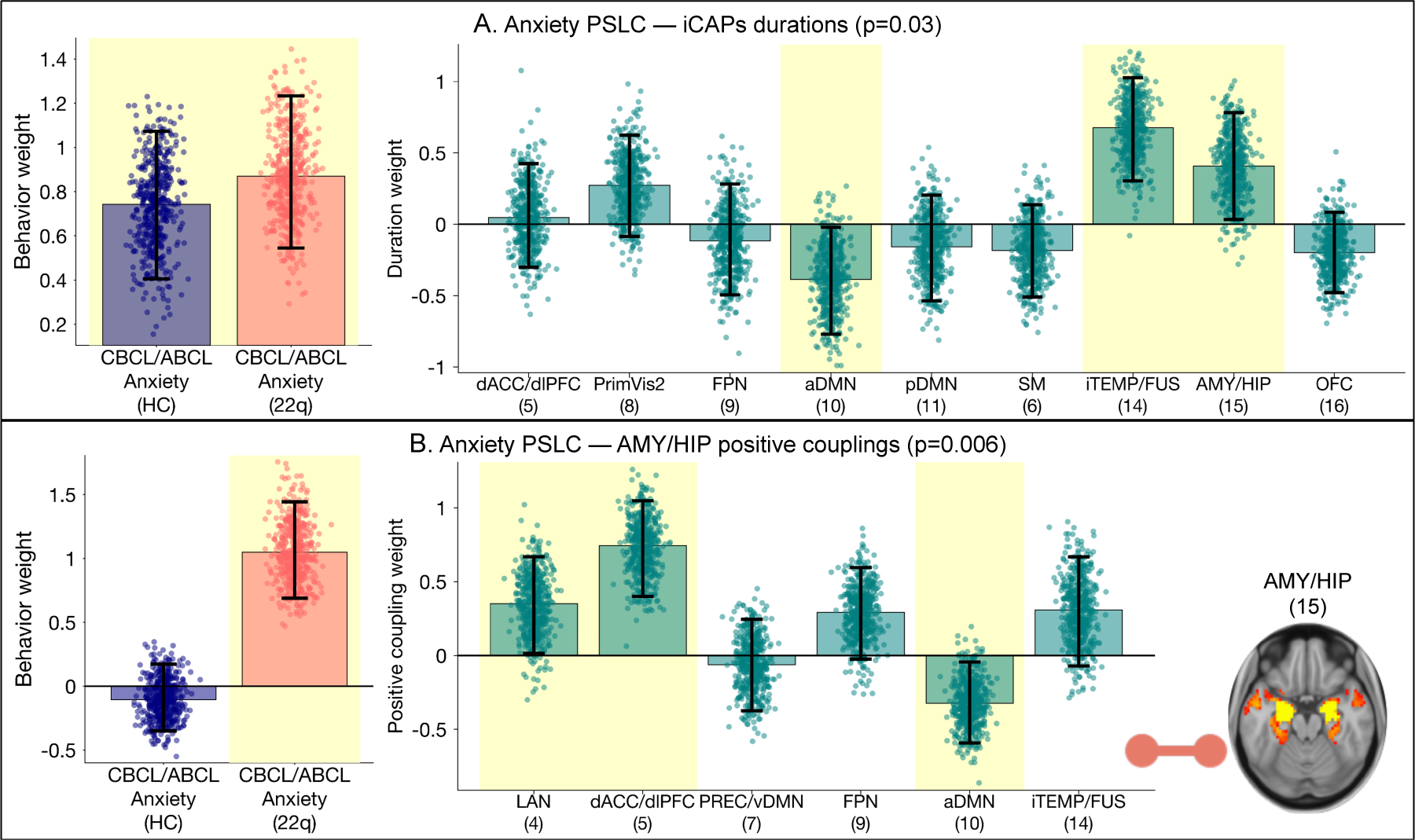
PLSC results for anxiety scores. A) Behavior weights and brain weights for PLSC including duration of nine altered iCAPs. There is a positive correlation of anxiety with duration of iTEMP/FUS (14) and AMY/HIP (15) and a negative correlation with duration of aDMN (10). B) Behavior weights and brain weights for PLSC including positive couplings of AMY/HIP (15). *Longer* positive coupling of AMY/HIP (15) with LAN (4) and dACC/dlPFC (5), and *shorter* positive coupling with aDMN (10) are associated with *higher* anxiety *only* in patients with 22q11DS. Error bars indicate bootstrapping 5th to 95th percentiles, robust results were indicated by yellow background. Exact values of bootstrap mean and 5-95 percentiles are reported in Supplementary Table S6. PLSC results for anti-couplings were not significant (p=0.07) and are thus not reported here.

### Altered couplings of AMY/HIP are associated with anxiety in patients with 22q11DS

To further investigate coupling effects related to anxiety, we conducted PLSC for coupling time of AMY/HIP (15). We have chosen to focus on this network because its duration was related to anxiety in the previous analysis (see previous paragraph “Altered iCAPs’ duration is associated with anxiety in patients with 22q11DS and HCs”) and because of the involvement of these brain regions in anxiety, which is well established in the literature (22). We included coupling time of AMY/HIP (15) with iCAPs that had altered couplings (iCAPs 4, 5, 7 and 9; see figure 3) and with iCAPs whose duration was significantly associated with anxiety (iCAPs 10 and 14; see figure 5A).

In the first PLSC analysis, testing for anti-couplings between AMY/HIP (15) and these networks, there was no significant correlation component (p=0.07).

The second PLSC analysis included positive couplings between AMY/HIP (15) and these networks and resulted in one significant correlation component (p=0.006, see figure 5B). Behavior weights were only robust for patients with 22q11DS, indicating that the corresponding pattern of correlations given by the correlation weights was specific for patients. Longer positive coupling of AMY/HIP (15) with LAN (4) and dACC/dlPFC (5) was positively associated with anxiety, whereas positive coupling with aDMN (10) was negatively associated with anxiety. Again, these results indicate that not only the activation of AMY/HIP (15) is related to anxiety, but also its coupling with other networks.

## Discussion

In this study, we investigated dynamic features of network brain activity in patients with 22q11DS, with a particular focus on the identification of functional signatures of prodromal psychotic symptoms and anxiety, two behavioral risk factors for the transition to psychosis. To the best of our knowledge, this is the first study to investigate dynamics of large-scale functional brain networks in 22q11DS. We used iCAPs to go beyond static functional connectivity analysis and look into precise moments of brain network activation and interaction, which is particularly promising to provide more sensitive imaging markers in schizophrenia (10). We observed alterations of brain networks’ duration and couplings in 22q11DS and associations between these patterns of alterations with positive psychotic symptoms and anxiety. We found shorter activation of fronto-parietal network (FPN), salience network (SN) and default-mode network (DMN) and longer activation of inferior temporal and limbic networks, as well as globally increased segregation between networks. Functional signatures of prodromal psychotic symptoms uncovered a pattern of relevant activation duration and coupling of dorsal anterior cingulate and dorsolateral prefrontal cortices (dACC/dlPFC). Functional signatures of anxiety revealed an implication of amygdala and hippocampus (AMY/HIP) activation duration and coupling, suggesting an adversarial effect of coupling with different sub-divisions of the ACC and medial PFC (mPFC), either reinforcing or protecting from anxiety.

### Alterations in 22q11DS: implication of cognitive and emotional brain networks

Comparing individuals with 22q11DS to healthy controls (HCs) revealed a varied pattern of longer and shorter network activations, suggesting that they ‘over-engage’ certain brain states, while ‘under-engaging’ in others. In particular, patients with 22q11DS presented shorter activation of FPN, DMN and cingulo-prefrontal SN (i.e., dACC/dlPFC). According to the triple-network hypothesis, these three core brain networks play a central role for higher cognitive functions and conversely their dysfunctional interaction could account for several psychiatric symptoms (57). Specifically, the hypothesis stipulates that the dynamic shift between an internally-oriented default state and the externally-directed goal-directed fronto-parietal state is mediated by the activation of a third network responsible for attributing salience. Of note, while these dynamic interactions are critical for higher cognitive functioning, they even occur at rest (28). Still, static connectivity approaches are unable to fully capture the proposed dynamic nature of such interactions. Here we observe reduced activation of all three networks in 22q11DS, which could suggest a less efficient engagement in these basic brain dynamics that are crucial to engage and disengage in goal-directed behavior. One could speculate that the disruption of these core mechanisms may underlie broad impairments in higher cognitive function described both in 22q11DS and psychosis (56, 35). In turn, there was longer activation in networks comprising limbic regions including amygdala, medial temporal and orbitofrontal cortices. While the dichotomy between cognitive and emotional brain is arguably artificial, longer activation in regions highly involved in emotional processing such as amygdala and orbitofrontal cortex could reflect higher emotional load during scanning in patients with 22q11DS (47, 68). The pattern of activation showed a significant, but different, development with age in both subgroups. Whilst in HCs higher age was associated with increased duration in FPN and cingulo-prefrontal SN and reduced duration of limbic networks, the opposite pattern was observed in 22q11DS. These results suggest that the atypical activation pattern observed in 22q11DS emerges with age, in accordance with the neurodevelopmental model of schizophrenia (35, 73).

Besides duration of activation, the iCAPs approach allowed us to probe the pattern of coupling between different networks. The pattern of altered couplings in 22q11DS was characterized by predominantly longer anti-couplings, accounting for more than half (13/25) of the alterations. Longer anti-coupling is suggestive of increased segregation between brain networks and is in agreement with evidence of increased segregation and decreased integration of structural and functional brain networks in both 22q11DS and non-syndromic psychosis (64, 76, 77, 79, 91, 93). Network segregation is a central feature of brain function that is important for cognition and attention (97) and its alterations in 22q11DS may be reflective of cognitive disabilities on a more global level than the above mentioned alterations in triple network activation that concentrates on three core networks.

### Functional signature of psychosis prodrome: aberrant salience network duration and coupling

The presence of prodromal psychotic symptoms was associated with longer activation of inferior temporal and fusiform (iTEMP/FUS), dACC/dlPFC and FPN. Increased activation of inferior temporal and fusiform gyrus has been previously reported in schizophrenia in terms of relative cerebral blood flow (44, 51) and BOLD variability (49). Also in 22q11DS, we observed higher BOLD variability in inferior temporal and fusiform regions in a partially overlapping sample (102), suggesting that increased BOLD variability might reflect longer network activation. Further, prodromal psychotic symptoms were associated with longer activation of the dACC/dlPFC network. The dACC is considered a key node of the SN involved in attributing subjective salience to internally and externally generated events (57, 86). Aberrant salience attribution has been proposed to possibly account for several positive psychotic symptoms, including hallucinations and delusions (37). Also electroencephalogram (EEG) studies in psychosis and 22q11DS have consistently reported longer representation of the EEG topography that corresponds to SN (7, 74, 88, 89), which, together with our findings, supports that aberrant salience processing may be a key mechanism in the emergence of psychotic symptoms in 22q11DS.

However, whilst duration of both dACC/dlPFC and FPN was positively correlated with psychotic symptoms, it was reduced overall in 22q11DS compared to HCs. Converging evidence from both structural and functional MRI points towards altered connectivity of the ACC in individuals with 22q11DS and psychotic symptoms (20, 76, 80, 101), reviewed in (65). Hence we suspected that the quality of the activations; i.e., the coupling with other networks, might be relevant for higher psychotic symptoms. Indeed, the analysis of dACC/dlPFC couplings revealed a significant relationship between higher psychotic symptoms and anti-coupling with both FPN and iTEMP/FUS. Taken together, these results suggest that whilst activations of dACC/dlPFC and FPN occur less frequently in 22q11DS in general, they are more frequently anti-coupled with one another and with iTEMP/FUS in patients with higher psychotic symptoms. The triple-network model proposes that activation of the SN is instrumental in re-orienting attention by mediating the shifts between DMN and FPN (57). Our findings of longer anti-coupling between SN and FPN suggest that this functional role of the cingulo-prefrontal SN is disrupted in individuals with higher psychotic symptoms, possibly reflecting mechanisms that underlie atypical salience processing (37).

Altogether, the richness of our iCAPs approach permitted to characterize a pattern reflecting SN activations that contribute to the pathophysiology of psychotic symptoms, both in terms of duration and quality. Our findings support the key role of network dynamics in the ACC in increased psychosis vulnerability (65) and point towards disrupted triple network function centered on the SN, which might reflect aberrant salience processing in patients with psychotic symptoms (57, 37).

### Functional signature of anxiety: aberrant amygdala & hippocampus duration and coupling

For both HCs and patients with 22q11DS anxiety was associated with a pattern of longer activation of AMY/HIP, iTEMP/FUS and primary visual (PrimVIS2) and shorter activation of anterior DMN (aDMN). Evidence in animal models and humans has revealed a central role of the amygdala in fear exposure, anticipation and reaction (47, 90, 16, 52). Anxiety in humans is associated with hyper-reactivity of the amygdala to fearful stimuli (22, 25). Also in absence of stimulation, increased metabolic activity in amygdala, hippocampus and inferior temporal cortex was found in rhesus monkeys with anxious temperament (26, 27) and cerebral blood flow in amygdala and fusiform cortex has been associated with trait anxiety in humans (36). The iCAPs approach allowed us to quantify moments of network activation and confirmed that hyperactivity AMY/HIP and iTEMP/FUS at rest could indeed represent trait markers of anxiety in both HCs and 22q11DS. Hyperactivity of both AMY/HIP and iTEMP/FUS observed in 22q11DS could therefore account for increased prevalence of anxiety disorders in this population.

Importantly, the amygdala does not operate in isolation, but is part of a complex circuit involved in regulating emotional responses (21). Indeed, in accord with the role in salience processing mentioned above, the dorsal portion of the ACC and PFC promote amygdala activity and are critical in the appraisal and expression of fear behavior (21). Oppositely, the subgenual-ACC and ventral mPFC largely dampen amygdala activity and are essential for fear extinction (21). This functional sub-division of the frontal lobe is also supported by extensive literature on fear circuitry in rodents, where the dorsal pre-limbic and the ventral infra-limbic cortices are found to have opposing roles on amygdala activation, fear expression, respectively fear extinction (90, 21, 58, 72, 94, 13, 14, 9). Given these findings, we speculated that the modulation of AMY/HIP activity particularly by the dACC/dlPFC and aDMN network may play a crucial role in the pathophysiology of anxiety. Indeed, we showed a significant positive association between anxiety and coupling duration between AMY/HIP and dACC/dlPFC, along with language network (LAN), FPN and iTEMP/FUS. Coupling duration between AMY/HIP and aDMN had an opposite, protective role on anxiety in accordance with the modulating role of mPFC-AMY projections on fear expression. Of note, the effects of amygdala coupling on anxiety appeared specific to individuals with 22q11DS, which could suggest that effects of amygdala modulation are nonlinear and relate only to more severe anxiety observed in 22q11DS.

In conclusion, we observed a dynamic functional pattern characterized both by longer AMY/HIP activations and atypical prefrontal AMY/HIP modulation, which might constitute a trait maker of anxiety and contribute vulnerability to psychosis in 22q11DS.

### Methodological aspects

#### The iCAPs framework

The present study is one of the first to apply the iCAPs framework in a clinical population and, due to the flexibility of the framework, we were able to discover distinct patterns of functional activation and interaction characteristic for prodromal psychotic symptoms and anxiety. The framework is unique in its ability to detect spatially *and* temporally overlapping networks (41, 103), and the robustness and richness of the presented results underlines its potential. Of note, extracted spatial patterns were highly similar to previously observed iCAPs retrieved from healthy subjects (41, 103), which reassures the performance of the iCAPs approach in a clinical population. Furthermore, the sub-division of classical resting-state networks such as the DMN and SN into multiple subnetworks confirms previously observed findings (41) and suggests that different subnetworks have distinct dynamic properties, which are difficult to detect by static approaches.

While iCAPs themselves were retrieved from a purely dynamic measure (i.e., the innovations), the measure of coupling duration between networks is closely linked to static functional connectivity (sFC). A direct comparison between the two showed that, as expected, alterations in couplings are related to aberrant sFC (see Supplementary Results and Supplementary Figure S6). However, while the results were similar, they were not identical, which is most likely due to the deconvolution, denoising and reconstruction of block-like time courses conducted before computing coupling duration. No such direct comparison was possible for activation duration, as it is a measure specific to each network which cannot be explained in terms of static connectivity.

#### BOLD signal analysis and motion

As in any fMRI study, we cannot exclude that the measured effects may have been influenced by non-neural confounds, such as heartbeat, respiration and motion (70). For the current study, we addressed this issue by taking typical measures during preprocessing, such as regression of WM and CSF signals, as well as motion censoring. Further, it is of note that the regularized deconvolution from the hemodynamic response function (HRF) further minimizes such effects, as any non-neural signals, which do not follow the HRF, are filtered out by the approach. Finally, we included group-centered average framewise displacement as nuisance regressor in all analyses to further avoid any confounding effects caused by motion in particular. To explicitly investigate whether observed group differences may have been caused by motion artifacts, we compared iCAPs activation duration in HCs with high motion versus low motion (see Supplementary Figure S5 and Supplementary Table S7). There were no significant differences in activation duration between these two groups, which suggests that motion was indeed not a major confound in our analysis. In patients with 22q11DS scanner motion is strongly correlated with the severity of symptoms, which makes it impossible to disentangle the two effects. Therefore, this remains a limitation of our study, despite the multiple measures we took to account for non-neural confounds.

## Conclusion

In summary, we here presented functional signatures of anxiety and positive psychotic symptoms in 22q11DS in terms of brain network activation and coupling. Our results confirm the implication of SN activity and connectivity in the emergence of psychotic symptoms. We further uncovered differential roles of dorsal and ventral ACC and mPFC coupling with AMY that are relevant for anxiety. Together, these findings shed light into the pathophysiology of two clinical risk factors that might represent relevant imaging markers for psychosis vulnerability.

## Methods and Materials

### Participants

The study included 221 subjects (111 patients with 22q11DS, 110 healthy controls (HCs), both aged 8–30 years). We excluded 33 patients and 25 HCs to ensure good data quality (see Supplementary Methods). The final sample included 78 patients with 22q11DS (37 males) and 85 HCs (36 males, see Table 1). HCs were recruited among patients’ siblings and through the Geneva state school system and had no present or past history of neurological or psychiatric disorders.

Prodromal positive psychotic symptoms in patients with 22q11DS were assessed using the Structured Interview of Prodromal Symptoms (SIPS; 61). The SIPS was not conducted in HCs. Anxiety was assessed both in HCs and patients with 22q11DS by combining the Child Behavioral Checklist (CBCL) Anxious-Depressed scale (1) in children, and the Adult Behavioral Checklist (ABCL) Anxious scale in adults above 18 years old (2).

Participants and their parents (for minors) gave their written informed consent and the research protocols were approved by the Institutional Review Board of Geneva University School of Medicine.

### Image acquisition

All MRI brain scans were acquired at the Centre d’Imagerie BioMédicale (CIBM) in Geneva on a Siemens Trio (12-channel coil; 54 HCs, 42 patients) and a Siemens Prisma (20-channel coil; 31 HCs, 36 patients) 3 Tesla scanner. Structural images were obtained with a T1-weighted sequence of 0.86*×*0.86*×*1.1 mm^3^ volumetric resolution (192 coronal slices, TR = 2500 ms, TE = 3 ms, acquisition matrix = 224 *×* 256, field of view = 22 cm^2^, flip angle = 8^*◦*^). Eyes-open rs-fMRI data were recorded with a T2*-weighted sequence of 8 minutes (voxel size = 1.84*×*1.84*×*3.2 mm, 38 axial slices, TR = 2400 ms, TE = 30 ms, flip angle = 85^*◦*^). Subjects were instructed to fixate a cross on the screen, let their mind wander and not to fall asleep.

### Preprocessing

Before applying the iCAPs pipeline, MRI scans were preprocessed using Statistical Parametric Mapping (SPM12, http://www.fil.ion.ucl.ac.uk/spm/) and functions of the Data Processing Assistant for Resting-State fMRI (DPARSF; 98) and Individual Brain Atlases using Statistical Parametric Mapping (IBASPM; 3) toolboxes. After realignment of functional scans, we applied spatial smoothing with an isotropic Gaussian kernel of 6 mm full width half maximum and coregistered structural scans to the functional mean. Structural images were segmented with the SPM12 *Segmentation* algorithm (5) and a study-specific template was generated using Diffeomorphic Anatomical Registration using Exponential Lie algebra (DARTEL; 4). Then, the first five functional scans were excluded and average white-matter and CSF signals were regressed out from the BOLD timeseries. We applied motion scrubbing (69) for the extended correction of motion artifacts, marking frames with a framewise displacement of more than 0.5 mm. As the filters implemented in the iCAPs framework require a constant sampling rate, marked frames were replaced by the spline interpolation of previous and following frames. Finally, motion frames were excluded before computation of temporal characteristics (described below).

### Total Activation and iCAPs

We used openly available Matlab code (https://c4science.ch/source/iCAPs/) to apply innovation-driven co-activation patterns (iCAPs; 41, 42, 103). We first employed Total Activation (TA; 39, 40, 23), which applies hemodynamically-informed deconvolution to the fMRI timeseries through spatio-temporal regularization. Significant activation changepoints (i.e., transients), derived from deconvolved timeseries, were concatenated across all subjects and fed into temporal K-means clustering to obtain simultaneously transitioning brain patterns, the iCAPs. The optimum number of clusters was determined by consensus clustering (62). Finally, time courses were obtained for all iCAPs using spatio-temporal transient-informed regression (103). A detailed description of all steps can be found in the Supplementary Methods.

### Extraction of temporal properties

For computation of temporal properties, iCAPs time courses were z-scored within each subject and thresholded at a z-score *> |*1*|* to determine ‘active’ timepoints (41). For each iCAP, we then computed the total duration of overall activation as percentage of the total non-motion scanning time.

Further, coupling and anti-coupling duration of two iCAPs were calculated as timepoints of same-signed or oppositely-signed co-activation measured as percentage of the total non-motion scanning time or as Jaccard score; i.e., percent joint activation time of the two respective iCAPs.

### Statistical analysis

#### Group comparisons of duration and coupling time

Duration and coupling measures between groups were compared using two-sample t-tests. P-values were corrected for multiple comparisons with the false discovery rate (FDR).

#### Partial Least Squares Correlation for multivariate relationship between iCAPs measures and behavioral variables

To evaluate multivariate patterns of correlation between behavioral variables and iCAPs activation measures, we used behavior partial least squares correlation (PLSC; 46). Briefly, we first computed a correlation matrix between behavioral variables (age, SIPS items, or CBCL/ABCL anxiety scores) and brain variables (iCAPs activation, positive coupling or anti-coupling duration). Group-specific correlation matrices of HCs and patients with 22q11DS were concatenated and singular value decomposition of this matrix then lead to several correlation components. Each correlation component is composed of a set of “behavior weights” and “iCAPs duration/coupling weights”. These weights indicate how strongly each variable contributes to the multivariate brain-behavior correlation in the associated correlation component. The weights lie between –1 and 1 and can be interpreted similarly to correlation values. Significance of correlation components was determined by permutation testing (1000 permutations) and stability of brain and behavior weights was obtained using bootstrapping (500 bootstrap samples). See Supplementary Methods for a detailed outline of PLSC.

We conducted multiple PLSC to investigate how psychotic symptoms and anxiety are related to the iCAPs’ activation and coupling measures. First, we conducted two PLSC analysis with the durations of altered iCAPs as brain measures and psychotic symptoms, respectively anxiety as behavioral measures. Then, we probed into behavioral effects of positive couplings and anti-couplings of one selected iCAP for each of the two behavioral measures, resulting in four more PLSC analyses. Due to the difference in design of each single PLSC in terms of type of measure and number of items, we did not correct for multiple comparisons.

#### Nuisance variable regression

Age, gender and motion were included as nuisance regressors in group comparisons and PLSC analyses. Nuisance regressors were standardized within each group to avoid linear dependence with the effects of interest.

## Acknowledgements

We are grateful to the subjects who participated in our study and thank Sarah Menghetti, Léa Chambaz, Virginie Pouillard and Dr. Maude Schneider for their involvement with the participants. We would also like to acknowledge Prof. François Lazeyras and the CIBM group for their support during data collection.

This work was supported by the Swiss National Science Foundation (SNSF) under Grants #32473B 121966, #234730 144260 and #145250 to S. Eliez, and Grant #163859 to M. Schaer, and by the National Center of Competence in Research (NCCR) “SYNAPSY – The Synaptic Bases of Mental Diseases”, SNSF, under Grants #51AU40 125759, #51NF40 158776 and #51NF40-185897.

