## Supplementary Material for "Large-scale brain network dynamics provide a measure of psychosis and anxiety in 22q11.2 deletion syndrome"

**Supplementary Results** **5**

**Supplementary Figures** **6**

**Supplementary Tables** **10**

**Supplementary References** **23**

*Corresponding author

*Preprint submitted to* *February 15, 2019*

### Supplementary Methods

*Participants Exclusion Criteria*

From the initial sample of 221 subjects between 8 and 30 years old, we had to exclude 33 patients and 25 controls to ensure good data quality. One control and one patient were excluded from the study because they reported having fallen asleep during the resting-state scan. Another 4 subjects (3 patients) were excluded due to high motion in the anatomical T1-weighted scan, 23 subjects (15 patients) were excluded due to excessive motion of more than 3 mm in translation or 3° in rotation in the resting-state fMRI scans and scans of another 19 subjects (7 patients) were not used because part of the cortex were not captured. Finally, 10 more subjects (7 patients) were excluded from the remaining sample after motion scrubbing (11) because less than 5 min of scanning time was remaining after censoring of frames with a framewise displacement higher than 0.5 mm. Motion data of the two groups is summarized in Supplementary Table S1.

*Overlapping Samples*

The cohort is partly overlapping with our previous resting-state fMRI studies: 31 subjects (16 patients) have been also included in Debbané et al. (1), 57 subjects (24 patients) in Scariati et al. (13), 62 subjects (27 patients) in Padula et al. (10), 59 subjects (26 patients) in Scariati et al. (12), 94 subjects (41 patients) in Padula et al. (9), 91 subjects (43 patients) in Zöller et al. (17) and 111 subjects (47 patients) in Zöller et al. (16).

*Total Activation and iCAPs*

The Total Activation (TA) and iCAPs framework is based on the detection of significant changepoints in deconvolved fMRI time series. Matlab code for the application of the whole framework can be found at `https://www.c4science.ch/source/iCAPs`.

In the first step, TA was used for the regularized deconvolution of fMRI signals from the hemodynamic response function (HRF) (3, 4, 2). In TA, the fMRI signal $\mathbf{y}(v)$ at voxel $v$ is modeled as a block-like activity-inducing signal $\mathbf{a}(v)$ convolved with the HRF $\mathbf{h}$ and additive white Gaussian noise $\boldsymbol{\epsilon}(v)$:

$$\mathbf{y}(v) = (\mathbf{a} \star \mathbf{h})(v) + \boldsymbol{\epsilon} = \mathbf{x}(v) + \boldsymbol{\epsilon}$$

Then the activity-related signal matrix $\mathbf{X}$ can be recovered by spatio-temporal regularization as follows:

$$\tilde{\mathbf{X}} = \arg\min_{\mathbf{X}} \frac{1}{2} ||\mathbf{Y} - \mathbf{X}||_F^2 + \mathcal{R}_T(\mathbf{X}) + \mathcal{R}_S(\mathbf{X}),$$

with the temporal regularization term

$$\mathcal{R}_T(\mathbf{X}) \quad = \sum_{v=1}^{N_v} \lambda_T(v) \sum_{t=1}^{N_t} |\Delta_L\{\mathbf{x}(v,\cdot)\}[t]|,$$

and the spatial regularization term

$$\mathcal{R}_S(\mathbf{X}) \quad = \lambda_S \sum_{t=1}^{N_t} \sum_{v=1}^{N_v} \sqrt{\sum_{u\in\mathcal{S}(v)} \left(x(v,t) - x(u,t)\right)^2}.$$

Here, $||\cdot||_F^2$ designates the Frobenius norm, $N_v$ is the number of voxels, $N_t$ is the number of time points, $\lambda_T$ is the temporal regularization parameter, $\lambda_S$ is the spatial regularization parameter, $\Delta_L = \Delta_D H^{-1}$ is a differential operator combining derivative $\Delta_D$ and HRF deconvolution $H^{-1}$ and $\mathcal{S}(v)$ is the neighborhood of voxel $v$.

After this regularized deconvolution from the HRF, innovation signals $\mathbf{i}$ were computed as the temporal derivative of the activity-inducing signals $\mathbf{a}$ and significant innovations were determined by a two-step thresholding procedure (5). First, a surrogate innovation distribution was computed at every voxel, and innovations larger than 95 % and lower than 5 % of the surrogate distribution were considered significant. Second, an innovation frame was considered significant, if there was a significant innovation in at least 5 % of the gray matter voxels.

Then, K-mean clustering was applied to the significant innovation frames, resulting in K spatial maps, the iCAPs. We used consensus clustering (8) to determine the optimum number of clusters.

Finally, iCAPs time courses were computed by transient-informed spatio-temporal back-projection of the spatial maps onto the activity-inducing signals (18). As spatial dependence of the iCAPs maps is possible, transient-informed regression is necessary to robustly recover temporal overlap in the iCAPs time courses. In the back-projection, transients in each iCAP's time course are restricted to frames with an innovation pattern that is close to the iCAP's spatial pattern (i.e., the cosine distance $d(t)$ between the innovation frame at time $t$ and the iCAP's spatial map has to respect $d(t) \leq \xi d_{min}(t)$ with $d_{min}(t)$ being the minimum distance of that innovation frame to any of the K iCAPs). Then the activation segments $\beta$ between two iCAP transients can be found for all iCAPs simultaneously by the following linear regression

$$\tilde{\boldsymbol{\beta}} = \arg\min_{\boldsymbol{\beta}} ||\mathbf{A}_C - \mathbf{S}\boldsymbol{\beta}||^2,$$

with the spatio-temporally concatenated activity-inducing signals $\mathbf{A}_C$, and the design matrix $\mathbf{S} = [\mathbf{S_1}|...|\mathbf{S}_K]$

containing one regressor with the repeated spatial iCAP map per activation segment $\beta$. Based on the Bayesian Information Criterion, in our data the optimum tuning parameter was $\xi = 1.3$. For a more detailed discussion of this transient-informed regression scheme, please refer to (18).

*Partial Least Squares Correlation*

Partial Least Squares Correlation (PLSC) is a well-suited method for the investigation of multivariate relationships between behavioral measures and brain activation data (6, 7). Here, we used so-called behavior PLSC to probe into relationships between demographic or clinical variables (i.e., age or psychotic symptoms/anxiety) and temporal properties of iCAPs (i.e., activation duration and coupling time). In the following we outline the steps of PLSC analysis. Variable annotations are used according to (6). For a more detailed discussion of PLSC analysis, we refer to (6, 7).

The first step in PLSC is the computation of group-wise correlation matrices $\mathbf{R}_{\mathrm{HC}} = \mathbf{Y}_{\mathrm{HC}}^{\top}\mathbf{X}_{\mathrm{HC}} \in \mathbb{R}^{N_{behav} \times N_{iCAPs}}$ between brain network properties $\mathbf{X}_{\mathrm{HC}} \in \mathbb{R}^{N_{\mathrm{HC}} \times N_{iCAPs}}$ and behavioral variables $\mathbf{Y}_{\mathrm{HC}} \in \mathbb{R}^{N_{\mathrm{HC}} \times N_{behav}}$ of HCs and $\mathbf{R}_{22\mathrm{q}} = \mathbf{Y}_{22\mathrm{q}}^{\top}\mathbf{X}_{22\mathrm{q}} \in \mathbb{R}^{N_{behav} \times N_{iCAPs}}$ between brain network properties $\mathbf{X}_{22\mathrm{q}} \in \mathbb{R}^{N_{22\mathrm{q}} \times N_{iCAPs}}$ and behavioral variables $\mathbf{Y}_{22\mathrm{q}} \in \mathbb{R}^{N_{22\mathrm{q}} \times N_{behav}}$ of patients with 22q11DS. $N_{22\mathrm{q}}$ is the number of patients with 22q11DS, $N_{\mathrm{HC}}$ is the number of HCs, $N_{iCAPs}$ is the number of included iCAPs measures and $N_{behav}$ is the number of included behavioral variables. $\mathbf{X}_{22\mathrm{q}}$, $\mathbf{X}_{\mathrm{HC}}$, $\mathbf{Y}_{22\mathrm{q}}$ and $\mathbf{Y}_{\mathrm{HC}}$ were z-scored across subjects before calculation of $\mathbf{R}_{22\mathrm{q}}$ and $\mathbf{R}_{\mathrm{HC}}$.

The common correlation matrix $\mathbf{R}$ is then computed by concatenating $\mathbf{R}_{22\mathrm{q}}$ and $\mathbf{R}_{22\mathrm{q}}$:

$$\mathbf{R} = \begin{bmatrix} \mathbf{R}_{\mathrm{HC}} \\ \mathbf{R}_{22\mathrm{q}} \end{bmatrix} = \begin{bmatrix} \mathbf{Y}_{\mathrm{HC}}^{\top}\mathbf{X}_{\mathrm{HC}} \\ \mathbf{Y}_{22\mathrm{q}}^{\top}\mathbf{X}_{22\mathrm{q}} \end{bmatrix} \in \mathbb{R}^{2N_{behav} \times N_{iCAPs}}.$$

Finally, $\mathbf{R}$ is decomposed into $2N_{behav}$ latent variables, or "correlation components", using singular value composition $\mathbf{R} = \mathbf{USV}^{\top}$. Each correlation component has a singular value (on the diagonal of $\mathbf{S}$) that specifies the explained correlation, as well as $2N_{behav}$ behavior saliences, or "behavior weights" (columns of $\mathbf{U}$), and $N_{iCAPs}$ duration/coupling saliences, or "duration/coupling weights" (rows of $\mathbf{V}^{\top}$). Behavior and brain weights indicate how strongly each variable contributes to the multivatiate behavior-brain correlation in a certain correlation component. They lie between $-1$ and 1 and – because we normalized the data before computation of $\mathbf{R}_{\mathrm{HC}}$ and $\mathbf{R}_{22\mathrm{q}}$ – can be interpreted similarly to correlation values. From the brain weights in $\mathbf{V}$, so called "brain scores" can be computed by projecting every subject's brain activation data onto the respective brain weights with $\mathbf{L}_{\mathbf{X}} = \mathbf{XV}$.

We used permutation testing with 1000 permutations to evaluate if any of the correlation components was significant and bootstrapping with 500 bootstrap samples to evaluate the stability of the behavior and brain weights. Significant PLSC results are reported in terms of bootstrapping mean and standard deviations.

**Supplementary Results**

We conducted a static functional connectivity (sFC) analysis to compare our results on dynamic network activity with a conventional static approach. For the measure of activation duration, there was no possibility to obtain a direct comparison, as it is specific to each network separately. The measure of coupling duration between networks, however, should be closely linked to sFC. In order to compare our iCAPs coupling measure and sFC, we computed regional averages from the preprocessed BOLD signals within each iCAP's spatial map, thresholded at a spatial z-score of 2.3. We then computed partial correlations between all pairs of iCAPs and compared the correlations between patients with 22q11DS and HCs. The results are shown in figure S6B. We can observe that while there are similarities, the results are not identical, which is most likely due to the deconvolution, denoising and reconstruction of block-like time courses which was conducted before computing coupling durations. Further, it is of note that with conventional sFC, it is impossible to detect alterations in both coupling and anti-coupling (such as we observe for iCAPs 5 and 3), as the correlation per definition only goes in one direction.

### Supplementary Figures

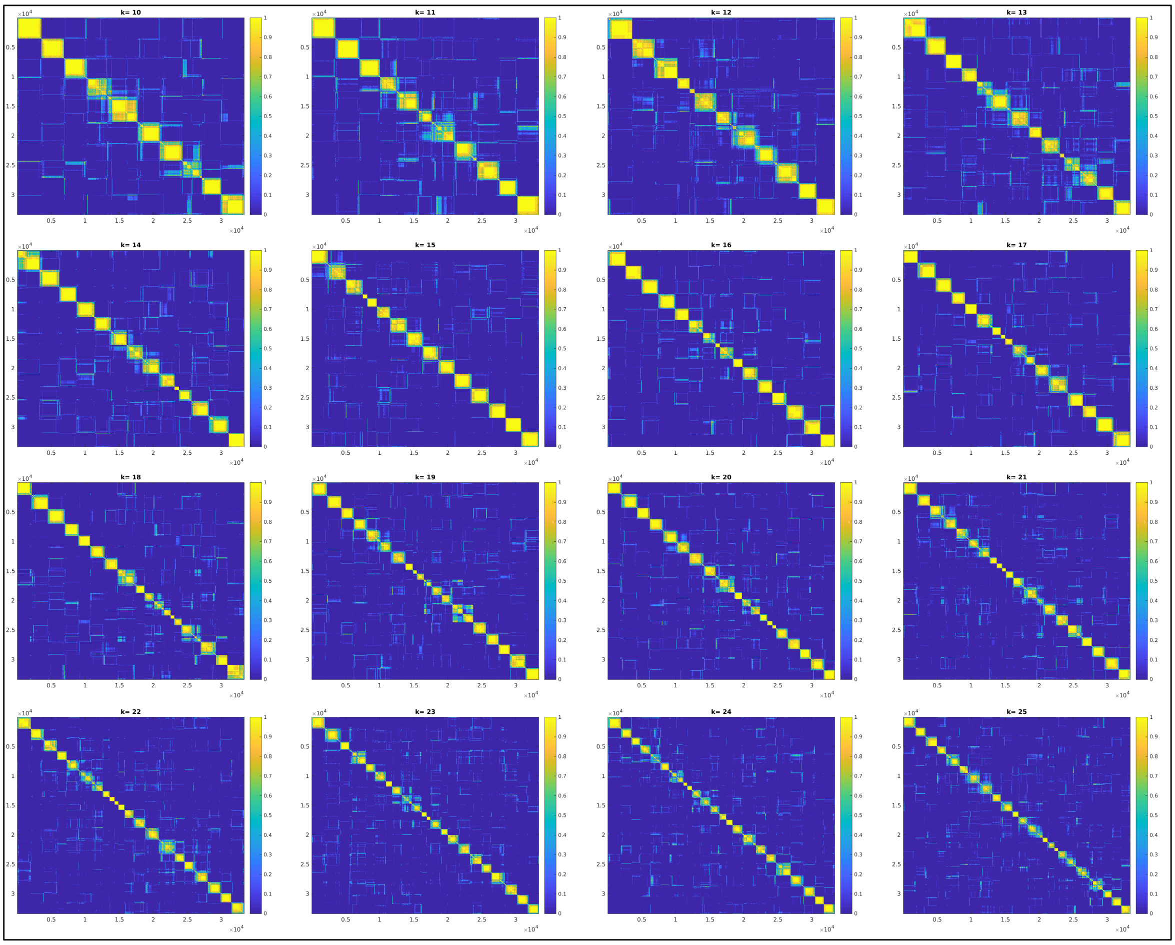

Figure S1: Consensus matrices for cluster numbers K=10 to K=25. The higher a value, the more often the two corresponding frames were clustered together during re-sampling. For optimum clustering, the consensus matrix should contain only zeros and ones (8). We selected to use $K = 17$ based on visual inspection of the consensus matrices and evaluation of consensus clustering quality measures (see Supplementary Figure S2).

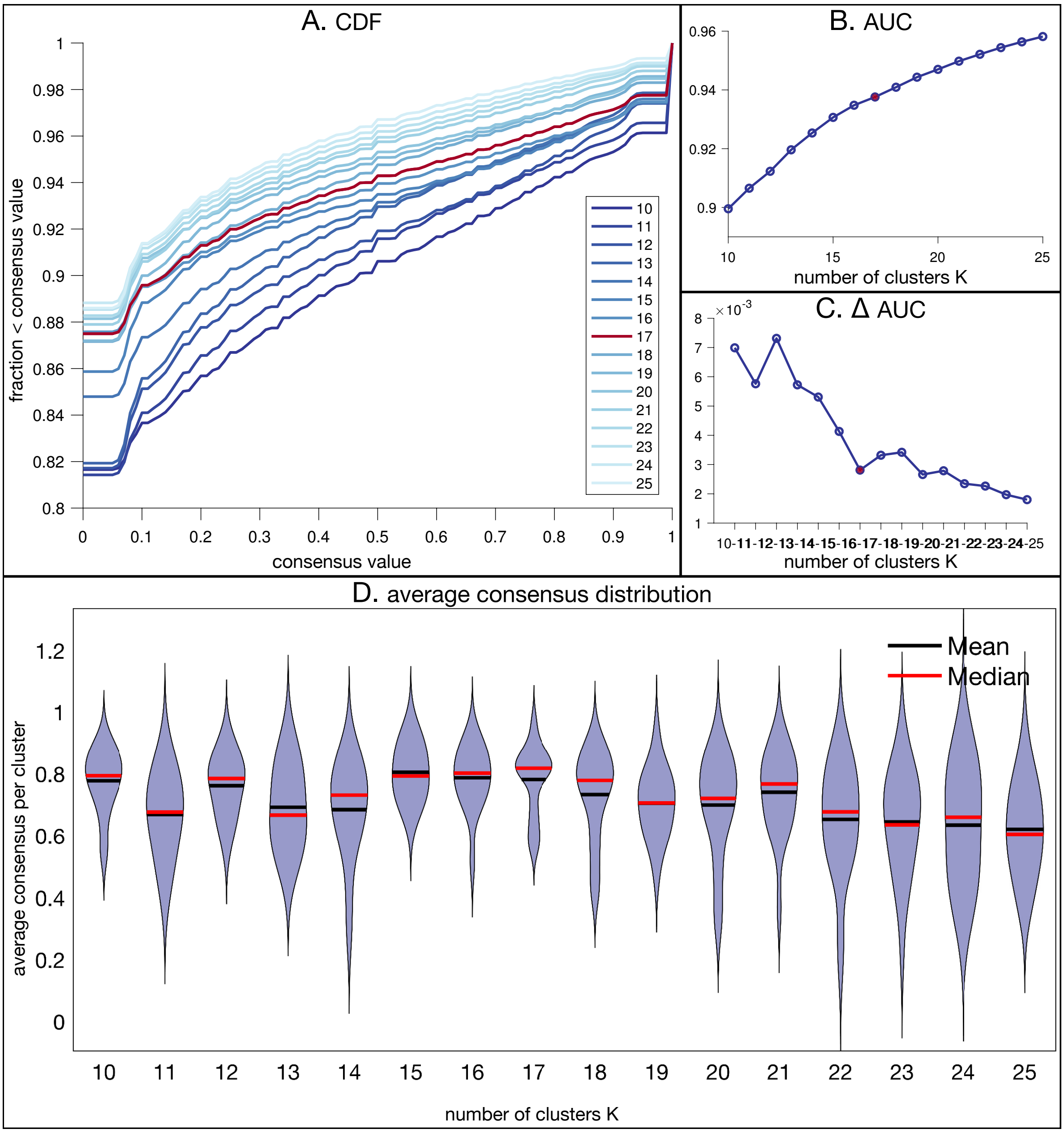

Figure S2: Consensus clustering quality measures for cluster numbers K=10 to K=25. A) the cumulative distribution function (CDF) of values in the respective consensus matrices (see Supplementary Figure S1). The optimum clustering solution would have a horizontal line with only values equal to 0 (two frames are never clustered together) or 1 (two frames are always clustered together). For K=17, the line is flatter than for surrounding clustering values. B) Area under the CDF curve and C) increase of the area under the curve with increasing K. D) Distribution of average consensus values per cluster. Median consensus is maximal for K=17. We selected to use $K = 17$ based on evaluation of the consensus clustering quality measures and visual inspection of the consensus matrices.

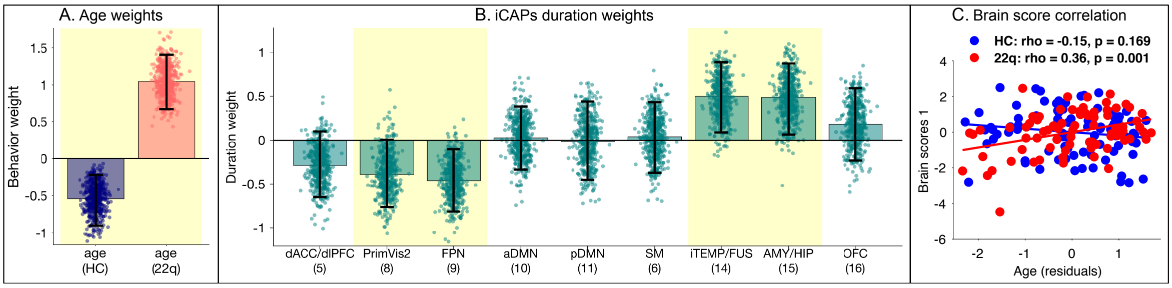

Figure S3: Age PLSC results (p=0.04). A) Age saliences indicating the strength and direction of age-relationship in HCs and patients with 22q11DS, respectively. B) iCAPs saliences indicating the strength and direction of each iCAP's duration-age-relationship. Activation of FPN (9) is shorter at higher age; activation of iTEMP/FUS (14) and AMY/HIP (15) is longer at higher age. Error bars indicate standard deviations of bootstrap distributions. C) Correlation of brain scores with age. As indicated by saliences in A, there is a positive correlation in patients with 22q11DS, and an opposite effect in HCs. Error bars indicate bootstrapping 5th to 95th percentiles, robust results were indicated by yellow background.

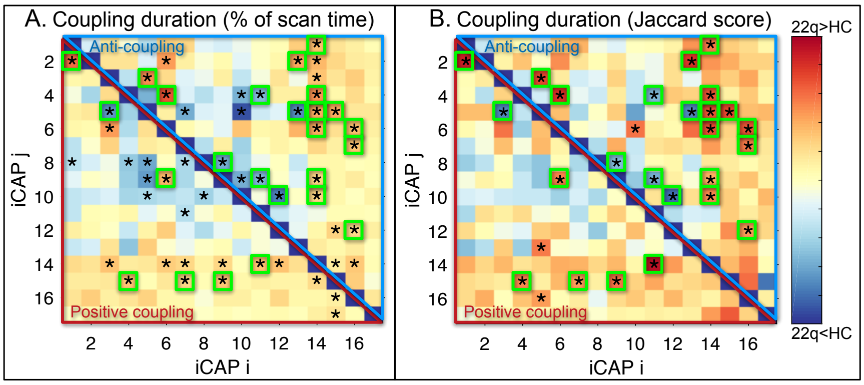

Figure S4: Duration differences of positive couplings (red triangles) and anti-couplings (blue triangles) between patients with 22q11DS and HCs. Corresponding test statistics (p-values, effect size) can be found in supplementary table S4. A) Coupling duration measured in percentage of total scanning time. B) Coupling duration measured in percentage of the joint activation time of the two respective iCAPs (Jaccard score). Significant differences (p<0.05) are marked with an asterisk (*), green boxes indicate coupling differences that are significant both in A and B. Couplings in green boxes were included in the graphical visualization in figure 3 of the main manuscript. P-values are FDR-corrected for multiple comparisons and age, gender and motion were included as covariates.

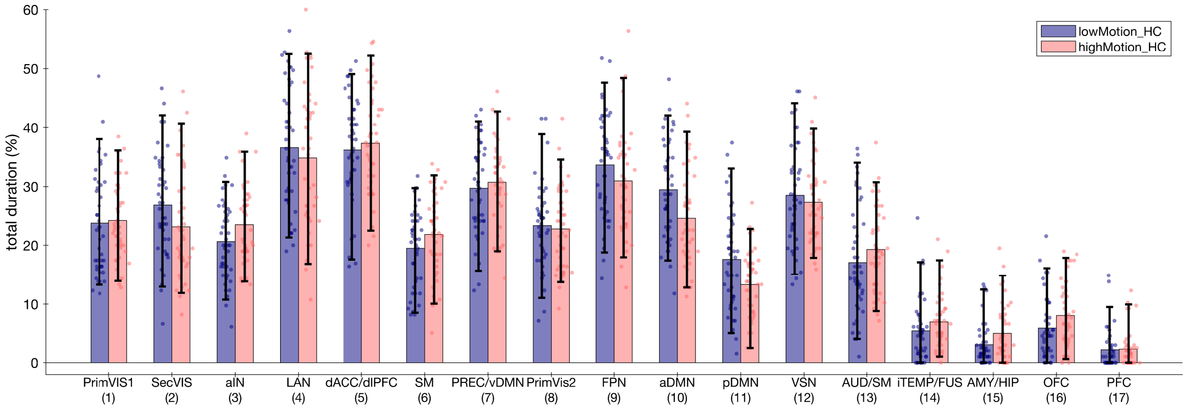

Figure S5: Group comparison of iCAPs' activation duration in high motion vs. low motion healthy subjects (divided by median split). There are no significant group differences between HCs with higher or lower motion. P-values are FDR-corrected for multiple comparisons and age, gender were included as covariates.

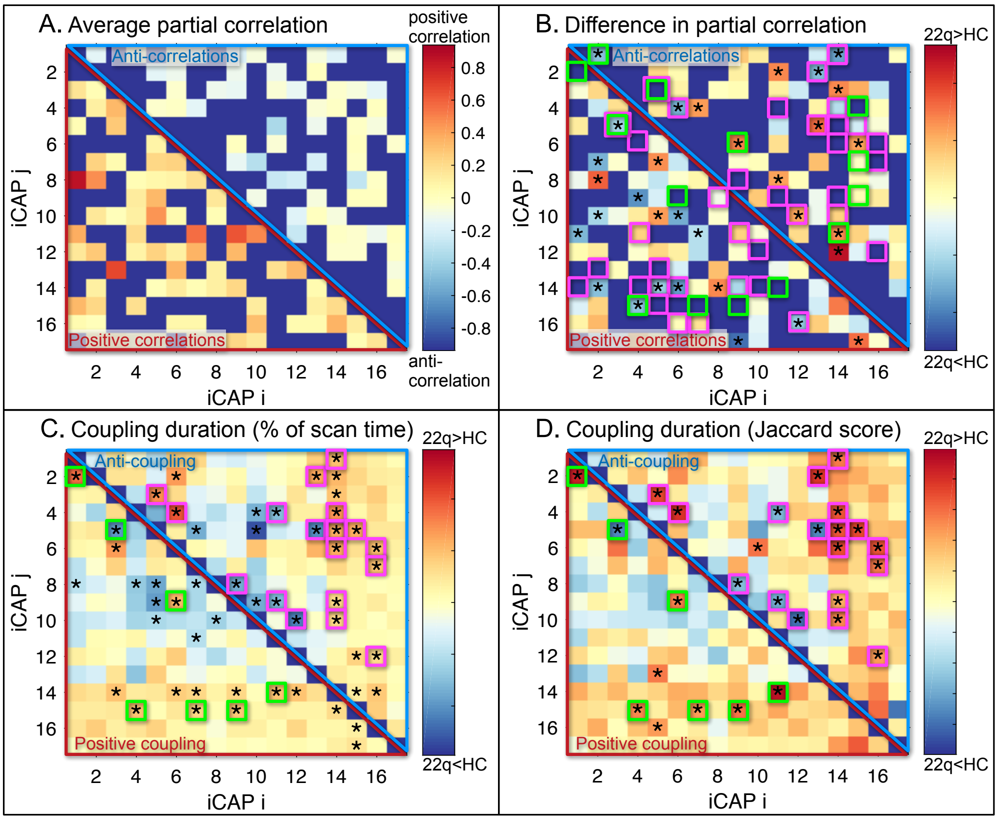

Figure S6: A) Average static functional connectivity (sFC) measured as partial correlations between average BOLD signals in each iCAP. The average was computed across all subjects including both patients with 22q11DS and healthy controls. B) Group differences in sFC between networks. C) Group differences in iCAPs coupling duration measured as percentage of total scan time. D) Group differences in iCAPs coupling duration measured as percentage of joint activation time (Jaccard score). Group differences in partial correlations are partially, but not entirely the same as group differences in positive couplings (green boxes) or anti-couplings (magenta boxes). The direction of alteration, if significant, is mostly the same in static FC and couplings; i.e., lower sFC corresponds to either lower positive coupling duration or higher anti-coupling (or both; e.g., iCAPs 5 and 3); higher sFC corresponds to either higher positive coupling of lower anti-coupling.

### Supplementary Tables

Table S1: fMRI motion data, FD - framewise displacement

| | **HC** | **22q11DS** | **p-value** |
|---|---|---|---|
| Mean FD (mm), before scrubbing | 0.15 ± 0.06 | 0.22 ± 0.10 | <0.001 |
| Mean FD (mm) after scrubbing | 0.13 ± 0.04 | 0.17 ± 0.06 | <0.001 |
| N. timepoints after scrubbing | 188.22 ± 12.87 | 178.26 ± 19.28 | <0.001 |

Table S2: iCAPs functional networks of Greicius atlas (14) and regions in the automated anatomical labeling (AAL) atlas (15). Percentiles indicate the fraction of voxels of a functional network or region that have a z-score > 2.3. A network/region is listed if more than 20% of the network/region is included in the iCAP.

| iCAP | Greicius network (%) | AAL Lobe | AAL Region (%) | z-score | voxels |
|---|---|---|---|---|---|
| 1 | Primary_Visual (81.12%) | Occipital | Calcarine_R (65.97%) | 3.76 | 221 |
| | | Occipital | Lingual_L (65.85%) | 3.62 | 295 |
| | | Occipital | Lingual_R (61.40%) | 3.73 | 272 |
| | | Occipital | Calcarine_L (50.45%) | 3.61 | 224 |
| | | Occipital | Cuneus_R (23.37%) | 2.98 | 61 |
| | | Occipital | Cuneus_L (22.86%) | 2.77 | 56 |
| 2 | Higher_Visual (75.94%) | Occipital | Occipital_Inf_L (80.47%) | 3.81 | 136 |
| | | Occipital | Occipital_Inf_R (77.33%) | 3.46 | 116 |
| | | Occipital | Occipital_Mid_R (65.61%) | 3.63 | 248 |
| | | Occipital | Occipital_Mid_L (56.79%) | 3.46 | 318 |
| | | Occipital | Fusiform_R (38.31%) | 3.25 | 218 |
| | | Occipital | Fusiform_L (36.94%) | 3.33 | 208 |
| | | Occipital | Occipital_Sup_R (32.39%) | 2.97 | 57 |
| | | Occipital | Occipital_Sup_L (22.42%) | 2.83 | 37 |
| | | Temporal | Temporal_Inf_R (20.19%) | 3.48 | 166 |
| 3 | Auditory (60.56%) | Temporal | Heschl_R (83.33%) | 2.88 | 45 |
| | Basal_Ganglia (20.13%) | Limbic | Insula_R (80.90%) | 3.58 | 288 |
| | | Central | Rolandic_Oper_L (77.55%) | 3.00 | 152 |
| | | Limbic | Insula_L (74.12%) | 3.35 | 335 |
| | | Central | Rolandic_Oper_R (71.79%) | 3.05 | 196 |
| | | Temporal | Heschl_L (71.67%) | 2.85 | 43 |
| | | Subcortical | Putamen_R (54.50%) | 2.91 | 103 |
| | | Subcortical | Putamen_L (50.00%) | 2.84 | 81 |
| | | Temporal | Temporal_Sup_L (38.04%) | 2.75 | 167 |
| | | Subcortical | Pallidum_L (25.00%) | 2.52 | 5 |
| | | Frontal | Frontal_Inf_Oper_L (23.56%) | 2.88 | 41 |
| | | Frontal | Frontal_Inf_Oper_R (21.24%) | 3.27 | 55 |
| 4 | Language (78.42%) | Temporal | Temporal_Mid_L (55.49%) | 3.33 | 551 |
| | | Temporal | Temporal_Mid_R (39.96%) | 3.13 | 356 |
| | | Parietal | Angular_L (39.27%) | 3.09 | 97 |
| | | Temporal | Temporal_Sup_R (36.31%) | 3.22 | 199 |
| 5 | Anterior_Salience (52.64%) | Frontal | Frontal_Mid_R (46.60%) | 3.27 | 432 |
| | | Frontal | Frontal_Mid_L (40.14%) | 3.18 | 334 |
| | | Limbic | Cingulum_Ant_R (28.25%) | 3.13 | 76 |
| | | Limbic | Cingulum_Ant_L (27.40%) | 3.12 | 77 |
| | | Limbic | Cingulum_Mid_R (21.86%) | 3.33 | 106 |
| 6 | Sensorimotor (53.69%) | Parietal | Paracentral_Lobule_L (74.15%) | 3.02 | 109 |
| | | Parietal | Paracentral_Lobule_R (60.19%) | 2.72 | 62 |
| | | Frontal | Supp_Motor_Area_R (59.60%) | 3.32 | 211 |
| | | Frontal | Supp_Motor_Area_L (51.11%) | 3.37 | 161 |
| | | Limbic | Cingulum_Mid_L (35.56%) | 3.08 | 144 |
| | | Parietal | Postcentral_R (35.53%) | 2.79 | 210 |
| | | Limbic | Cingulum_Mid_R (33.81%) | 3.06 | 164 |
| | | Frontal | Precentral_L (33.20%) | 3.07 | 172 |
| | | Frontal | Precentral_R (30.77%) | 2.93 | 156 |
| | | Parietal | Postcentral_L (30.69%) | 2.84 | 174 |

| iCAP | Greicius network (%) | AAL Lobe | AAL Region (%) | mean z-score | voxels |
|---|---|---|---|---|---|
| 7 | Precuneus (65.04%) | Parietal | Precuneus_R (61.35%) | 3.32 | 319 |
| | Ventral_DMN (38.00%) | Parietal | Precuneus_L (51.19%) | 3.39 | 280 |
| | | Occipital | Occipital_Sup_R (44.32%) | 3.19 | 78 |
| | | Parietal | Parietal_Sup_L (43.09%) | 3.11 | 134 |
| | | Parietel | Parietal_Sup_R (41.90%) | 3.07 | 106 |
| | | Occipital | Occipital_Sup_L (40.61%) | 3.09 | 67 |
| | | Occipital | Cuneus_R (33.33%) | 2.74 | 87 |
| | | Occipital | Occipital_Mid_R (28.31%) | 3.00 | 107 |
| | | Occipital | Cuneus_L (25.71%) | 2.62 | 63 |
| 8 | Primary_Visual (93.71%) | Occipital | Calcarine_R (70.15%) | 4.32 | 235 |
| | Higher_Visual (22.74%) | Occipital | Calcarine_L (65.32%) | 4.00 | 290 |
| | | Occipital | Cuneus_R (54.79%) | 3.33 | 143 |
| | | Occipital | Cuneus_L (54.29%) | 3.47 | 133 |
| | | Occipital | Lingual_R (51.47%) | 3.45 | 228 |
| | | Occipital | Lingual_L (49.78%) | 3.37 | 223 |
| | | Occipital | Occipital_Sup_R (44.32%) | 3.05 | 78 |
| | | Occipital | Occipital_Sup_L (40.61%) | 3.01 | 67 |
| 9 | Left_ECN (60.21%) | Parietal | Angular_L (64.37%) | 3.37 | 159 |
| | Right_ECN (47.88%) | Parietal | Angular_R (59.64%) | 3.47 | 201 |
| | Precuneus (21.68%) | Parietal | Parietal_Inf_R (42.44%) | 3.65 | 115 |
| | | Parietal | Parietal_Inf_L (34.55%) | 3.46 | 171 |
| | | Frontal | Frontal_Mid_L (27.40%) | 2.89 | 228 |
| 10 | Dorsal_DMN (58.75%) | Limbic | Cingulum_Ant_R (73.23%) | 3.93 | 197 |
| | | Frontal | Frontal_Sup_Orb_Medial_L (73.11%) | 4.17 | 87 |
| | | Limbic | Cingulum_Ant_L (69.75%) | 4.23 | 196 |
| | | Frontal | Frontal_Sup_Orb_Medial_R (69.48%) | 3.87 | 107 |
| | | Frontal | Frontal_Sup_Medial_L (54.95%) | 3.65 | 211 |
| | | Frontal | Frontal_Sup_Medial_R (46.30%) | 3.64 | 169 |
| | | Limbic | Cingulum_Post_L (35.00%) | 2.62 | 28 |
| | | Frontal | Rectus_L (34.68%) | 3.17 | 60 |
| | | Frontal | Rectus_R (28.47%) | 3.23 | 39 |
| | | Limbic | Cingulum_Post_R (25.00%) | 2.51 | 8 |
| 11 | Precuneus (34.73%) | Limbic | Cingulum_Post_L (97.50%) | 5.47 | 78 |
| | Dorsal_DMN (24.30%) | Limbic | Cingulum_Post_R (96.88%) | 5.55 | 31 |
| | | Parietal | Angular_L (61.94%) | 2.94 | 153 |
| | | Parietal | Precuneus_R (45.77%) | 4.24 | 238 |
| | | Parietal | Angular_R (43.92%) | 2.95 | 148 |
| | | Parietal | Precuneus_L (39.31%) | 4.47 | 215 |
| 12 | Visuospatial (61.54%) | Parietal | Parietal_Inf_R (60.89%) | 3.59 | 165 |
| | Posterior_Salience (26.50%) | Parietal | Parietal_Inf_L (56.77%) | 3.44 | 281 |
| | | Frontal | Frontal_Inf_Oper_R (43.24%) | 3.01 | 112 |
| | | Parietal | SupraMarginal_R (33.00%) | 3.33 | 133 |
| | | Parietal | Postcentral_R (24.37%) | 3.21 | 144 |
| | | Parietal | SupraMarginal_L (23.36%) | 3.20 | 57 |
| | | Frontal | Frontal_Inf_Oper_L (22.99%) | 2.96 | 40 |
| | | Frontal | Frontal_Inf_Tri_R (20.74%) | 2.56 | 67 |

| iCAP | Greicius network (%) | AAL Lobe | AAL Region (%) | mean z-score | voxels |
|---|---|---|---|---|---|
| 13 | Auditory (46.06%) | Central | Rolandic_Oper_R (76.56%) | 3.26 | 209 |
| | Sensorimotor (25.59%) | Central | Rolandic_Oper_L (63.78%) | 2.99 | 125 |
| | Posterior_Salience (20.35%) | Parietal | Postcentral_L (46.56%) | 3.85 | 264 |
| | | Parietal | Postcentral_R (37.23%) | 3.93 | 220 |
| | | Temporal | Heschl_R (37.04%) | 2.59 | 20 |
| | | Parietal | SupraMarginal_R (36.97%) | 3.22 | 149 |
| | | Parietal | SupraMarginal_L (34.02%) | 3.05 | 83 |
| | | Frontal | Precentral_R (24.46%) | 3.55 | 124 |
| 14 | | Temporal | Temporal_Inf_L (55.30%) | 3.87 | 386 |
| | | Temporal | Temporal_Inf_R (46.84%) | 3.87 | 385 |
| | | Occipital | Fusiform_R (22.67%) | 4.23 | 129 |
| 15 | | Limbic | Amygdala_L (82.76%) | 3.97 | 48 |
| | | Limbic | Amygdala_R (79.25%) | 3.54 | 42 |
| | | Temporal | Temporal_Pole_Mid_L (74.26%) | 3.30 | 101 |
| | | Limbic | Hippocampus_L (70.29%) | 4.56 | 123 |
| | | Temporal | Temporal_Pole_Mid_R (69.89%) | 2.95 | 130 |
| | | Limbic | ParaHippocampal_L (67.46%) | 3.67 | 114 |
| | | Limbic | ParaHippocampal_R (65.74%) | 3.75 | 142 |
| | | Limbic | Hippocampus_R (60.37%) | 4.36 | 99 |
| | | Occipital | Fusiform_L (30.37%) | 3.04 | 171 |
| | | Temporal | Temporal_Pole_Sup_L (28.07%) | 3.06 | 48 |
| | | Temporal | Temporal_Pole_Sup_R (22.91%) | 2.73 | 41 |
| | | Occipital | Fusiform_R (21.44%) | 2.87 | 122 |
| 16 | | Frontal | Rectus_R (90.51%) | 4.38 | 124 |
| | | Frontal | Rectus_L (83.82%) | 4.02 | 145 |
| | | Frontal | Olfactory_R (80.36%) | 3.99 | 45 |
| | | Frontal | Olfactory_L (76.62%) | 3.63 | 59 |
| | | Frontal | Frontal_Sup_Orb_R (53.37%) | 4.80 | 95 |
| | | Frontal | Frontal_Sup_Orb_L (50.00%) | 4.92 | 85 |
| | | Frontal | Frontal_Mid_Orb_L (37.71%) | 3.69 | 66 |
| | | Frontal | Frontal_Mid_Orb_R (30.93%) | 3.57 | 60 |
| | | Frontal | Frontal_Inf_Orb_L (25.96%) | 3.68 | 88 |
| | | Frontal | Frontal_Inf_Orb_R (21.64%) | 3.76 | 74 |
| 17 | | Frontal | Frontal_Mid_Orb_R (63.40%) | 5.61 | 123 |
| | | Frontal | Frontal_Mid_Orb_L (57.14%) | 5.21 | 100 |
| | | Frontal | Frontal_Sup_Orb_R (50.00%) | 6.55 | 89 |
| | | Frontal | Frontal_Sup_Orb_L (48.82%) | 7.10 | 83 |
| | | Frontal | Frontal_Sup_Orb_Medial_L (43.70%) | 4.89 | 52 |
| | | Frontal | Frontal_Sup_Orb_Medial_R (35.06%) | 5.91 | 54 |

Table S3: Test statistics corresponding to figure 2 of the main manuscript: Results from two-sample t-tests comparing each iCAP's activation duration between healthy controls and patients with 22q11DS. Age, gender and scanner motion (framewise displacement) were included as nuisance regressors. P-values were corrected for multiple comparisons using false discovery rate.

| iCAP | t-statistic | p-value | effect size (Cohen's d) |
|---|---|---|---|
| PrimVIS1 (1) | -0.16 | 0.875 | 0.02 |
| SecVIS (2) | 0.80 | 0.480 | -0.13 |
| aIN (3) | 1.24 | 0.285 | -0.19 |
| LAN (4) | -2.17 | 0.053 | 0.34 |
| dACC/dlPFC (5) | -3.88 | 0.001 | 0.61 |
| SM (6) | 2.60 | 0.019 | -0.41 |
| PREC/vDMN (7) | -1.50 | 0.209 | 0.24 |
| PrimVis2 (8) | -3.49 | 0.002 | 0.55 |
| FPN (9) | -3.25 | 0.004 | 0.51 |
| aDMN (10) | -4.78 | 0.000 | 0.75 |
| pDMN (11) | -3.03 | 0.006 | 0.48 |
| VSN (12) | -0.64 | 0.556 | 0.10 |
| AUD/SM (13) | -1.04 | 0.364 | 0.16 |
| iTEMP/FUS (14) | 4.98 | 0.000 | -0.78 |
| AMY/HIP (15) | 3.71 | 0.001 | -0.58 |
| OFC (16) | 3.20 | 0.004 | -0.50 |
| PFC (17) | 1.24 | 0.285 | -0.19 |

Table S4: Test statistics corresponding to figure 3 and Supplementary Figure S4: Results from two-sample t-tests comparing positive coupling time and anti-coupling time of pairwise iCAP combinations between healthy controls and patients with 22q11DS. Age, gender and scanner motion (framewise displacement) were included as nuisance regressors. P-values were corrected for multiple comparisons using false discovery rate.

| | | **Percentage of scan time** | | | **Jaccard score** | | |
|---|---|---|---|---|---|---|---|
| | iCAP | t-statistic | p-value | effect size (Cohen's d) | t-statistic | p-value | effect size (Cohen's d) |
| **positive coupling** | 2 - 1 | 3.79 | 0.005 | -0.59 | 4.18 | 0.002 | -0.65 |
| | 3 - 1 | 0.52 | 0.715 | -0.08 | 0.12 | 0.942 | -0.02 |
| | 4 - 1 | -0.64 | 0.661 | 0.10 | -0.01 | 0.997 | 0.00 |
| | 5 - 1 | 0.69 | 0.641 | -0.11 | 1.71 | 0.278 | -0.27 |
| | 6 - 1 | 0.39 | 0.788 | -0.06 | 0.10 | 0.944 | -0.02 |
| | 7 - 1 | -1.75 | 0.208 | 0.28 | -1.44 | 0.340 | 0.23 |
| | 8 - 1 | -2.81 | 0.036 | 0.44 | -2.62 | 0.072 | 0.41 |
| | 9 - 1 | -1.01 | 0.501 | 0.16 | -0.47 | 0.783 | 0.07 |
| | 10 - 1 | -1.77 | 0.203 | 0.28 | -1.00 | 0.501 | 0.16 |
| | 11 - 1 | -2.59 | 0.049 | 0.41 | -1.85 | 0.242 | 0.29 |
| | 12 - 1 | 0.73 | 0.630 | -0.11 | 1.04 | 0.494 | -0.16 |
| | 13 - 1 | -1.97 | 0.149 | 0.31 | -2.38 | 0.109 | 0.37 |
| | 14 - 1 | 2.18 | 0.110 | -0.34 | 2.32 | 0.120 | -0.36 |
| | 15 - 1 | 1.63 | 0.234 | -0.25 | 1.51 | 0.333 | -0.24 |
| | 16 - 1 | 1.66 | 0.233 | -0.26 | 1.00 | 0.501 | -0.16 |
| | 17 - 1 | -0.89 | 0.569 | 0.14 | -1.04 | 0.494 | 0.16 |
| | 3 - 2 | 0.66 | 0.658 | -0.10 | 0.71 | 0.642 | -0.11 |
| | 4 - 2 | -2.08 | 0.124 | 0.33 | -1.68 | 0.282 | 0.27 |
| | 5 - 2 | 0.96 | 0.530 | -0.15 | 1.78 | 0.263 | -0.28 |
| | 6 - 2 | 0.32 | 0.827 | -0.05 | -0.23 | 0.888 | 0.04 |
| | 7 - 2 | -0.13 | 0.923 | 0.02 | -0.37 | 0.810 | 0.06 |
| | 8 - 2 | -1.36 | 0.349 | 0.21 | -1.45 | 0.340 | 0.23 |
| | 9 - 2 | -0.15 | 0.907 | 0.02 | 0.45 | 0.788 | -0.07 |
| | 10 - 2 | -1.64 | 0.234 | 0.26 | -1.02 | 0.497 | 0.16 |
| | 11 - 2 | 0.30 | 0.836 | -0.05 | 1.02 | 0.497 | -0.16 |
| | 12 - 2 | 0.26 | 0.840 | -0.04 | 0.10 | 0.944 | -0.02 |
| | 13 - 2 | -1.64 | 0.234 | 0.26 | -1.73 | 0.277 | 0.27 |
| | 14 - 2 | 2.18 | 0.110 | -0.34 | 1.49 | 0.340 | -0.23 |
| | 15 - 2 | 1.91 | 0.164 | -0.30 | 1.31 | 0.386 | -0.21 |
| | 16 - 2 | 2.61 | 0.048 | -0.41 | 2.13 | 0.168 | -0.33 |
| | 17 - 2 | 0.89 | 0.568 | -0.14 | 0.44 | 0.788 | -0.07 |
| | 4 - 3 | 1.29 | 0.385 | -0.20 | 1.32 | 0.383 | -0.21 |
| | 5 - 3 | -4.33 | 0.002 | 0.68 | -3.99 | 0.003 | 0.63 |
| | 6 - 3 | 2.96 | 0.031 | -0.46 | 2.46 | 0.093 | -0.39 |
| | 7 - 3 | 1.81 | 0.194 | -0.28 | 1.96 | 0.211 | -0.31 |
| | 8 - 3 | -0.94 | 0.542 | 0.15 | -0.23 | 0.888 | 0.04 |
| | 9 - 3 | -0.54 | 0.705 | 0.08 | 0.03 | 0.987 | -0.00 |
| | 10 - 3 | -1.62 | 0.241 | 0.25 | -1.47 | 0.340 | 0.23 |
| | 11 - 3 | 0.07 | 0.958 | -0.01 | 0.70 | 0.642 | -0.11 |
| | 12 - 3 | -0.29 | 0.839 | 0.05 | -0.64 | 0.684 | 0.10 |
| | 13 - 3 | 1.60 | 0.246 | -0.25 | 1.47 | 0.340 | -0.23 |
| | 14 - 3 | 3.19 | 0.018 | -0.50 | 1.78 | 0.263 | -0.28 |
| | 15 - 3 | 1.92 | 0.162 | -0.30 | 0.86 | 0.576 | -0.14 |
| | 16 - 3 | 1.18 | 0.423 | -0.18 | 0.00 | 0.997 | -0.00 |
| | 17 - 3 | 0.60 | 0.675 | -0.10 | 0.46 | 0.787 | -0.07 |

| | iCAP | Percentage of scan time t-statistic | p-value | effect size (Cohen's d) | Jaccard score t-statistic | p-value | effect size (Cohen's d) |
|---|---|---|---|---|---|---|---|
| **positive coupling** | 5 - 4 | -2.22 | 0.107 | 0.35 | -1.44 | 0.340 | 0.23 |
| | 6 - 4 | -2.00 | 0.142 | 0.31 | -1.93 | 0.220 | 0.30 |
| | 7 - 4 | -1.18 | 0.423 | 0.19 | -0.42 | 0.801 | 0.07 |
| | 8 - 4 | -2.82 | 0.036 | 0.44 | -1.99 | 0.202 | 0.31 |
| | 9 - 4 | -2.39 | 0.074 | 0.38 | -1.43 | 0.340 | 0.22 |
| | 10 - 4 | -2.24 | 0.103 | 0.35 | -1.31 | 0.386 | 0.21 |
| | 11 - 4 | 0.54 | 0.705 | -0.09 | 1.42 | 0.342 | -0.22 |
| | 12 - 4 | -1.73 | 0.214 | 0.27 | -1.04 | 0.494 | 0.16 |
| | 13 - 4 | -2.60 | 0.049 | 0.41 | -2.06 | 0.186 | 0.32 |
| | 14 - 4 | 1.27 | 0.388 | -0.20 | 1.11 | 0.457 | -0.17 |
| | 15 - 4 | 3.37 | 0.012 | -0.52 | 3.65 | 0.007 | -0.57 |
| | 16 - 4 | 1.39 | 0.338 | -0.22 | 1.81 | 0.253 | -0.28 |
| | 17 - 4 | 0.86 | 0.573 | -0.13 | 1.34 | 0.377 | -0.21 |
| | 6 - 5 | 1.19 | 0.423 | -0.19 | 1.58 | 0.305 | -0.25 |
| | 7 - 5 | 0.63 | 0.661 | -0.10 | 1.56 | 0.310 | -0.24 |
| | 8 - 5 | -3.52 | 0.010 | 0.55 | -2.51 | 0.088 | 0.39 |
| | 9 - 5 | -3.60 | 0.008 | 0.57 | -2.62 | 0.072 | 0.41 |
| | 10 - 5 | -2.77 | 0.037 | 0.44 | -1.73 | 0.277 | 0.27 |
| | 11 - 5 | -1.59 | 0.252 | 0.25 | -0.62 | 0.692 | 0.10 |
| | 12 - 5 | 0.58 | 0.691 | -0.09 | 1.39 | 0.350 | -0.22 |
| | 13 - 5 | 2.04 | 0.131 | -0.32 | 3.09 | 0.030 | -0.48 |
| | 14 - 5 | 1.28 | 0.385 | -0.20 | 1.58 | 0.305 | -0.25 |
| | 15 - 5 | 0.45 | 0.757 | -0.07 | 0.89 | 0.562 | -0.14 |
| | 16 - 5 | 2.38 | 0.075 | -0.37 | 3.08 | 0.030 | -0.48 |
| | 17 - 5 | 0.28 | 0.839 | -0.04 | 0.40 | 0.809 | -0.06 |
| | 7 - 6 | 0.63 | 0.661 | -0.10 | 0.53 | 0.766 | -0.08 |
| | 8 - 6 | -0.04 | 0.977 | 0.01 | 0.12 | 0.942 | -0.02 |
| | 9 - 6 | 2.77 | 0.037 | -0.43 | 3.35 | 0.018 | -0.52 |
| | 10 - 6 | -2.21 | 0.107 | 0.35 | -1.99 | 0.202 | 0.31 |
| | 11 - 6 | -0.39 | 0.788 | 0.06 | -0.26 | 0.876 | 0.04 |
| | 12 - 6 | 1.91 | 0.163 | -0.30 | 1.71 | 0.278 | -0.27 |
| | 13 - 6 | -0.37 | 0.801 | 0.06 | -0.23 | 0.888 | 0.04 |
| | 14 - 6 | 2.92 | 0.033 | -0.45 | 2.10 | 0.170 | -0.33 |
| | 15 - 6 | 2.01 | 0.139 | -0.31 | 1.98 | 0.203 | -0.31 |
| | 16 - 6 | 0.78 | 0.603 | -0.12 | 0.16 | 0.941 | -0.02 |
| | 17 - 6 | 1.22 | 0.409 | -0.19 | 0.91 | 0.553 | -0.14 |
| | 8 - 7 | -2.73 | 0.040 | 0.43 | -2.11 | 0.170 | 0.33 |
| | 9 - 7 | -2.53 | 0.056 | 0.40 | -1.47 | 0.340 | 0.23 |
| | 10 - 7 | -2.20 | 0.107 | 0.35 | -1.43 | 0.340 | 0.22 |
| | 11 - 7 | -2.92 | 0.033 | 0.46 | -2.10 | 0.170 | 0.33 |
| | 12 - 7 | -1.11 | 0.460 | 0.17 | -0.73 | 0.637 | 0.11 |
| | 13 - 7 | -1.14 | 0.446 | 0.18 | -0.50 | 0.774 | 0.08 |
| | 14 - 7 | 2.69 | 0.042 | -0.42 | 2.76 | 0.056 | -0.43 |
| | 15 - 7 | 4.01 | 0.004 | -0.62 | 4.40 | 0.002 | -0.68 |
| | 16 - 7 | 0.86 | 0.573 | -0.14 | 1.02 | 0.497 | -0.16 |
| | 17 - 7 | 1.21 | 0.416 | -0.19 | 1.58 | 0.305 | -0.25 |

| | iCAP | Percentage of scan time t-statistic | p-value | effect size (Cohen's d) | Jaccard score t-statistic | p-value | effect size (Cohen's d) |
|---|---|---|---|---|---|---|---|
| **positive coupling** | 9 - 8 | -0.69 | 0.641 | 0.11 | 0.47 | 0.783 | -0.07 |
| | 10 - 8 | -3.39 | 0.012 | 0.53 | -2.26 | 0.135 | 0.35 |
| | 11 - 8 | -0.87 | 0.573 | 0.14 | 0.12 | 0.942 | -0.02 |
| | 12 - 8 | -2.66 | 0.044 | 0.42 | -2.39 | 0.109 | 0.38 |
| | 13 - 8 | -0.27 | 0.839 | 0.04 | 0.14 | 0.941 | -0.02 |
| | 14 - 8 | 1.23 | 0.408 | -0.19 | 1.29 | 0.388 | -0.20 |
| | 15 - 8 | 0.54 | 0.705 | -0.08 | 0.75 | 0.630 | -0.12 |
| | 16 - 8 | -1.24 | 0.407 | 0.19 | -1.25 | 0.401 | 0.20 |
| | 17 - 8 | -0.48 | 0.742 | 0.07 | -0.38 | 0.810 | 0.06 |
| | 10 - 9 | -1.58 | 0.252 | 0.25 | -0.09 | 0.944 | 0.01 |
| | 11 - 9 | 0.68 | 0.641 | -0.11 | 1.69 | 0.280 | -0.26 |
| | 12 - 9 | -1.18 | 0.423 | 0.19 | -0.39 | 0.810 | 0.06 |
| | 13 - 9 | 0.26 | 0.840 | -0.04 | 1.58 | 0.305 | -0.25 |
| | 14 - 9 | 2.90 | 0.033 | -0.45 | 2.67 | 0.064 | -0.42 |
| | 15 - 9 | 3.15 | 0.019 | -0.49 | 3.40 | 0.017 | -0.53 |
| | 16 - 9 | 1.41 | 0.328 | -0.22 | 1.64 | 0.293 | -0.26 |
| | 17 - 9 | 0.65 | 0.659 | -0.10 | 1.12 | 0.457 | -0.17 |
| | 11 - 10 | -1.49 | 0.298 | 0.23 | -0.04 | 0.986 | 0.01 |
| | 12 - 10 | 0.50 | 0.730 | -0.08 | 1.69 | 0.280 | -0.26 |
| | 13 - 10 | -2.13 | 0.115 | 0.34 | -0.70 | 0.642 | 0.11 |
| | 14 - 10 | 0.18 | 0.889 | -0.03 | 0.46 | 0.787 | -0.07 |
| | 15 - 10 | 0.74 | 0.625 | -0.12 | 1.66 | 0.288 | -0.26 |
| | 16 - 10 | 0.97 | 0.530 | -0.15 | 1.27 | 0.393 | -0.20 |
| | 17 - 10 | -0.28 | 0.839 | 0.04 | 0.60 | 0.712 | -0.09 |
| | 12 - 11 | -0.68 | 0.641 | 0.11 | -0.32 | 0.840 | 0.05 |
| | 13 - 11 | -0.77 | 0.604 | 0.12 | -0.15 | 0.941 | 0.02 |
| | 14 - 11 | 4.44 | 0.002 | -0.69 | 4.78 | 0.001 | -0.74 |
| | 15 - 11 | 1.78 | 0.202 | -0.28 | 2.29 | 0.128 | -0.35 |
| | 16 - 11 | 0.29 | 0.839 | -0.05 | 0.71 | 0.638 | -0.11 |
| | 17 - 11 | 0.80 | 0.590 | -0.12 | 0.74 | 0.637 | -0.12 |
| | 13 - 12 | -0.93 | 0.544 | 0.15 | -1.11 | 0.457 | 0.17 |
| | 14 - 12 | 3.38 | 0.012 | -0.53 | 2.82 | 0.049 | -0.44 |
| | 15 - 12 | 1.74 | 0.208 | -0.27 | 1.83 | 0.250 | -0.29 |
| | 16 - 12 | 2.04 | 0.131 | -0.32 | 2.03 | 0.195 | -0.32 |
| | 17 - 12 | 0.82 | 0.583 | -0.13 | 0.91 | 0.552 | -0.14 |
| | 14 - 13 | 2.18 | 0.110 | -0.34 | 0.77 | 0.629 | -0.12 |
| | 15 - 13 | 1.41 | 0.328 | -0.22 | 1.35 | 0.374 | -0.21 |
| | 16 - 13 | 0.72 | 0.633 | -0.11 | 0.86 | 0.576 | -0.14 |
| | 17 - 13 | 0.00 | 0.997 | -0.00 | 0.76 | 0.630 | -0.12 |
| | 15 - 14 | 2.87 | 0.034 | -0.44 | 1.16 | 0.437 | -0.18 |
| | 16 - 14 | 1.17 | 0.424 | -0.18 | 0.58 | 0.718 | -0.09 |
| | 17 - 14 | 2.11 | 0.118 | -0.33 | 2.00 | 0.202 | -0.31 |
| | 16 - 15 | 2.71 | 0.041 | -0.42 | 1.20 | 0.418 | -0.19 |
| | 17 - 15 | 2.90 | 0.033 | -0.45 | 2.47 | 0.093 | -0.38 |
| | 17 - 16 | 1.64 | 0.234 | -0.26 | 1.51 | 0.333 | -0.24 |

| | | Percentage of scan time | | | Jaccard score | | |
|---|---|---|---|---|---|---|---|
| | iCAP | t-statistic | p-value | effect size (Cohen's d) | t-statistic | p-value | effect size (Cohen's d) |
| **anti-coupling** | 1 - 2 | -1.28 | 0.385 | 0.20 | -1.31 | 0.386 | 0.21 |
| | 1 - 3 | 0.70 | 0.641 | -0.11 | 0.48 | 0.783 | -0.08 |
| | 1 - 4 | -0.33 | 0.827 | 0.05 | 0.10 | 0.944 | -0.02 |
| | 1 - 5 | -1.79 | 0.200 | 0.28 | -1.00 | 0.501 | 0.16 |
| | 1 - 6 | 1.64 | 0.234 | -0.26 | 1.29 | 0.388 | -0.20 |
| | 1 - 7 | 1.67 | 0.228 | -0.26 | 2.22 | 0.145 | -0.35 |
| | 1 - 8 | -0.85 | 0.573 | 0.13 | -0.43 | 0.798 | 0.07 |
| | 1 - 9 | -1.54 | 0.270 | 0.24 | -1.08 | 0.473 | 0.17 |
| | 1 - 10 | -1.09 | 0.468 | 0.17 | -0.13 | 0.942 | 0.02 |
| | 1 - 11 | -0.23 | 0.860 | 0.04 | 0.02 | 0.992 | -0.00 |
| | 1 - 12 | 0.24 | 0.849 | -0.04 | 0.48 | 0.783 | -0.07 |
| | 1 - 13 | 2.17 | 0.111 | -0.34 | 2.59 | 0.074 | -0.41 |
| | 1 - 14 | 3.40 | 0.012 | -0.53 | 3.27 | 0.020 | -0.51 |
| | 1 - 15 | 1.75 | 0.208 | -0.27 | 1.39 | 0.350 | -0.22 |
| | 1 - 16 | 1.37 | 0.344 | -0.21 | 1.17 | 0.433 | -0.18 |
| | 1 - 17 | 0.51 | 0.723 | -0.08 | 0.40 | 0.810 | -0.06 |
| | 2 - 3 | 1.03 | 0.496 | -0.16 | 0.38 | 0.810 | -0.06 |
| | 2 - 4 | 0.81 | 0.588 | -0.13 | 1.44 | 0.340 | -0.23 |
| | 2 - 5 | -0.94 | 0.542 | 0.15 | -0.14 | 0.942 | 0.02 |
| | 2 - 6 | 3.18 | 0.018 | -0.50 | 2.53 | 0.085 | -0.40 |
| | 2 - 7 | 0.57 | 0.692 | -0.09 | 0.79 | 0.621 | -0.12 |
| | 2 - 8 | -0.41 | 0.777 | 0.06 | 0.10 | 0.944 | -0.02 |
| | 2 - 9 | 0.41 | 0.777 | -0.07 | 0.94 | 0.536 | -0.15 |
| | 2 - 10 | -0.95 | 0.540 | 0.15 | -0.34 | 0.826 | 0.05 |
| | 2 - 11 | -0.28 | 0.839 | 0.04 | -0.77 | 0.629 | 0.12 |
| | 2 - 12 | 0.48 | 0.742 | -0.07 | 0.91 | 0.552 | -0.14 |
| | 2 - 13 | 3.20 | 0.018 | -0.50 | 3.65 | 0.007 | -0.57 |
| | 2 - 14 | 3.45 | 0.011 | -0.54 | 2.52 | 0.085 | -0.39 |
| | 2 - 15 | 2.51 | 0.059 | -0.39 | 1.89 | 0.228 | -0.30 |
| | 2 - 16 | 1.86 | 0.179 | -0.29 | 1.71 | 0.278 | -0.27 |
| | 2 - 17 | 2.37 | 0.077 | -0.37 | 2.19 | 0.149 | -0.34 |
| | 3 - 4 | -1.44 | 0.319 | 0.23 | -1.09 | 0.468 | 0.17 |
| | 3 - 5 | 2.90 | 0.033 | -0.45 | 3.69 | 0.007 | -0.57 |
| | 3 - 6 | 1.43 | 0.325 | -0.22 | 0.72 | 0.637 | -0.11 |
| | 3 - 7 | -1.13 | 0.450 | 0.18 | -1.21 | 0.413 | 0.19 |
| | 3 - 8 | -0.90 | 0.562 | 0.14 | -0.86 | 0.576 | 0.14 |
| | 3 - 9 | 0.85 | 0.573 | -0.13 | 1.48 | 0.340 | -0.23 |
| | 3 - 10 | -1.94 | 0.157 | 0.31 | -1.45 | 0.340 | 0.23 |
| | 3 - 11 | -0.84 | 0.575 | 0.13 | -0.76 | 0.630 | 0.12 |
| | 3 - 12 | 1.35 | 0.352 | -0.21 | 1.16 | 0.437 | -0.18 |
| | 3 - 13 | -0.80 | 0.590 | 0.12 | -0.79 | 0.621 | 0.12 |
| | 3 - 14 | 3.13 | 0.020 | -0.49 | 2.43 | 0.100 | -0.38 |
| | 3 - 15 | 2.21 | 0.107 | -0.34 | 1.68 | 0.282 | -0.26 |
| | 3 - 16 | 2.48 | 0.062 | -0.39 | 2.19 | 0.149 | -0.34 |
| | 3 - 17 | 1.67 | 0.228 | -0.26 | 1.16 | 0.437 | -0.18 |

| | iCAP | Percentage of scan time<br>t-statistic | <br>p-value | <br>effect size (Cohen's d) | Jaccard score<br>t-statistic | <br>p-value | <br>effect size (Cohen's d) |
|---|---|---|---|---|---|---|---|
| **anti-** | 4 - 5 | -2.08 | 0.125 | 0.33 | -1.00 | 0.501 | 0.16 |
| **coupling** | 4 - 6 | 3.96 | 0.004 | -0.62 | 4.14 | 0.002 | -0.64 |
| | 4 - 7 | -1.05 | 0.488 | 0.16 | -0.73 | 0.637 | 0.11 |
| | 4 - 8 | -2.07 | 0.127 | 0.33 | -1.40 | 0.347 | 0.22 |
| | 4 - 9 | -0.87 | 0.573 | 0.14 | 0.45 | 0.788 | -0.07 |
| | 4 - 10 | -2.82 | 0.036 | 0.45 | -1.29 | 0.388 | 0.20 |
| | 4 - 11 | -3.67 | 0.007 | 0.58 | -3.02 | 0.033 | 0.48 |
| | 4 - 12 | 0.57 | 0.691 | -0.09 | 1.42 | 0.340 | -0.22 |
| | 4 - 13 | 0.68 | 0.641 | -0.11 | 1.29 | 0.388 | -0.20 |
| | 4 - 14 | 4.96 | 0.000 | -0.77 | 4.65 | 0.001 | -0.72 |
| | 4 - 15 | 0.76 | 0.609 | -0.12 | 0.72 | 0.637 | -0.11 |
| | 4 - 16 | 1.94 | 0.157 | -0.30 | 1.80 | 0.260 | -0.28 |
| | 4 - 17 | 0.63 | 0.661 | -0.10 | 0.49 | 0.781 | -0.08 |
| | 5 - 6 | 0.54 | 0.705 | -0.09 | 1.27 | 0.393 | -0.20 |
| | 5 - 7 | -2.81 | 0.036 | 0.44 | -1.65 | 0.288 | 0.26 |
| | 5 - 8 | -1.19 | 0.423 | 0.19 | -0.15 | 0.941 | 0.02 |
| | 5 - 9 | -0.64 | 0.661 | 0.10 | 1.22 | 0.410 | -0.19 |
| | 5 - 10 | -3.35 | 0.012 | 0.53 | -2.19 | 0.149 | 0.34 |
| | 5 - 11 | -1.75 | 0.208 | 0.28 | -0.49 | 0.780 | 0.08 |
| | 5 - 12 | -0.97 | 0.530 | 0.15 | 0.27 | 0.868 | -0.04 |
| | 5 - 13 | -3.79 | 0.005 | 0.60 | -3.17 | 0.025 | 0.50 |
| | 5 - 14 | 3.56 | 0.009 | -0.55 | 4.11 | 0.002 | -0.64 |
| | 5 - 15 | 3.28 | 0.014 | -0.51 | 3.75 | 0.007 | -0.58 |
| | 5 - 16 | 1.23 | 0.408 | -0.19 | 1.92 | 0.221 | -0.30 |
| | 5 - 17 | 1.69 | 0.224 | -0.26 | 2.33 | 0.120 | -0.36 |
| | 6 - 7 | 0.33 | 0.827 | -0.05 | -0.10 | 0.944 | 0.02 |
| | 6 - 8 | -0.44 | 0.765 | 0.07 | -0.51 | 0.769 | 0.08 |
| | 6 - 9 | -1.08 | 0.471 | 0.17 | -0.94 | 0.536 | 0.15 |
| | 6 - 10 | 2.44 | 0.068 | -0.38 | 3.30 | 0.020 | -0.51 |
| | 6 - 11 | 0.68 | 0.641 | -0.11 | 0.67 | 0.664 | -0.11 |
| | 6 - 12 | 0.71 | 0.635 | -0.11 | 0.35 | 0.821 | -0.06 |
| | 6 - 13 | 2.66 | 0.044 | -0.41 | 2.72 | 0.060 | -0.42 |
| | 6 - 14 | 4.54 | 0.001 | -0.71 | 3.77 | 0.007 | -0.59 |
| | 6 - 15 | 2.50 | 0.060 | -0.39 | 1.39 | 0.350 | -0.22 |
| | 6 - 16 | 4.10 | 0.003 | -0.64 | 4.20 | 0.002 | -0.66 |
| | 6 - 17 | 2.13 | 0.115 | -0.33 | 1.90 | 0.225 | -0.30 |
| | 7 - 8 | -2.15 | 0.112 | 0.34 | -1.85 | 0.242 | 0.29 |
| | 7 - 9 | 0.63 | 0.661 | -0.10 | 1.81 | 0.253 | -0.28 |
| | 7 - 10 | -2.15 | 0.112 | 0.34 | -1.25 | 0.401 | 0.20 |
| | 7 - 11 | 0.06 | 0.963 | -0.01 | 0.95 | 0.530 | -0.15 |
| | 7 - 12 | 0.46 | 0.749 | -0.07 | 0.85 | 0.582 | -0.13 |
| | 7 - 13 | 0.28 | 0.839 | -0.04 | 0.85 | 0.582 | -0.13 |
| | 7 - 14 | 2.35 | 0.079 | -0.37 | 2.71 | 0.060 | -0.42 |
| | 7 - 15 | 1.41 | 0.328 | -0.22 | 1.63 | 0.296 | -0.25 |
| | 7 - 16 | 2.84 | 0.035 | -0.44 | 3.20 | 0.024 | -0.50 |
| | 7 - 17 | 1.28 | 0.385 | -0.20 | 1.47 | 0.340 | -0.23 |

| | iCAP | Percentage of scan time t-statistic | p-value | effect size (Cohen's d) | Jaccard score t-statistic | p-value | effect size (Cohen's d) |
|---|---|---|---|---|---|---|---|
| **anti-coupling** | 8 - 9 | -3.82 | 0.005 | 0.60 | -2.97 | 0.035 | 0.47 |
| | 8 - 10 | -2.43 | 0.068 | 0.38 | -1.24 | 0.401 | 0.20 |
| | 8 - 11 | -1.39 | 0.337 | 0.22 | 0.20 | 0.913 | -0.03 |
| | 8 - 12 | 0.02 | 0.990 | -0.00 | 0.42 | 0.798 | -0.07 |
| | 8 - 13 | -2.49 | 0.060 | 0.39 | -1.62 | 0.297 | 0.26 |
| | 8 - 14 | 1.04 | 0.491 | -0.16 | 1.04 | 0.494 | -0.16 |
| | 8 - 15 | 0.86 | 0.573 | -0.13 | 0.81 | 0.609 | -0.13 |
| | 8 - 16 | 0.26 | 0.840 | -0.04 | 0.38 | 0.810 | -0.06 |
| | 8 - 17 | 1.05 | 0.490 | -0.16 | 1.61 | 0.298 | -0.25 |
| | 9 - 10 | -4.13 | 0.003 | 0.65 | -2.74 | 0.058 | 0.43 |
| | 9 - 11 | -3.89 | 0.004 | 0.61 | -3.01 | 0.033 | 0.47 |
| | 9 - 12 | -0.02 | 0.990 | 0.00 | 0.64 | 0.684 | -0.10 |
| | 9 - 13 | -1.45 | 0.319 | 0.23 | -0.52 | 0.769 | 0.08 |
| | 9 - 14 | 2.77 | 0.037 | -0.43 | 2.89 | 0.042 | -0.45 |
| | 9 - 15 | 1.02 | 0.501 | -0.16 | 1.46 | 0.340 | -0.23 |
| | 9 - 16 | 1.09 | 0.470 | -0.17 | 1.72 | 0.278 | -0.27 |
| | 9 - 17 | 0.13 | 0.923 | -0.02 | 0.73 | 0.637 | -0.11 |
| | 10 - 11 | -0.85 | 0.573 | 0.13 | 0.73 | 0.637 | -0.11 |
| | 10 - 12 | -3.37 | 0.012 | 0.53 | -3.27 | 0.020 | 0.51 |
| | 10 - 13 | -0.32 | 0.829 | 0.05 | 1.00 | 0.501 | -0.16 |
| | 10 - 14 | 2.68 | 0.042 | -0.42 | 2.97 | 0.035 | -0.47 |
| | 10 - 15 | 0.85 | 0.573 | -0.13 | 1.66 | 0.288 | -0.26 |
| | 10 - 16 | -0.10 | 0.942 | 0.02 | 0.37 | 0.810 | -0.06 |
| | 10 - 17 | 1.04 | 0.490 | -0.16 | 2.02 | 0.199 | -0.31 |
| | 11 - 12 | -1.87 | 0.178 | 0.29 | -1.50 | 0.336 | 0.23 |
| | 11 - 13 | -1.84 | 0.185 | 0.29 | -1.26 | 0.401 | 0.20 |
| | 11 - 14 | -1.34 | 0.352 | 0.21 | -1.23 | 0.403 | 0.19 |
| | 11 - 15 | 0.59 | 0.683 | -0.09 | 1.59 | 0.305 | -0.25 |
| | 11 - 16 | 0.36 | 0.804 | -0.06 | 0.75 | 0.630 | -0.12 |
| | 11 - 17 | 0.41 | 0.778 | -0.06 | 1.06 | 0.486 | -0.17 |
| | 12 - 13 | -0.83 | 0.578 | 0.13 | -0.31 | 0.846 | 0.05 |
| | 12 - 14 | 0.81 | 0.587 | -0.13 | 0.39 | 0.810 | -0.06 |
| | 12 - 15 | 2.77 | 0.037 | -0.43 | 2.87 | 0.043 | -0.45 |
| | 12 - 16 | 2.87 | 0.034 | -0.45 | 3.07 | 0.030 | -0.48 |
| | 12 - 17 | 2.14 | 0.114 | -0.33 | 2.32 | 0.120 | -0.36 |
| | 13 - 14 | 1.72 | 0.214 | -0.27 | 1.12 | 0.457 | -0.18 |
| | 13 - 15 | 0.80 | 0.590 | -0.13 | 1.24 | 0.402 | -0.19 |
| | 13 - 16 | 1.80 | 0.197 | -0.28 | 1.73 | 0.277 | -0.27 |
| | 13 - 17 | 0.19 | 0.882 | -0.03 | -0.19 | 0.913 | 0.03 |
| | 14 - 15 | 2.69 | 0.042 | -0.42 | 1.77 | 0.267 | -0.28 |
| | 14 - 16 | 3.00 | 0.029 | -0.47 | 1.44 | 0.340 | -0.23 |
| | 14 - 17 | 1.06 | 0.484 | -0.17 | 0.29 | 0.860 | -0.05 |
| | 15 - 16 | 2.62 | 0.048 | -0.41 | 1.93 | 0.220 | -0.30 |
| | 15 - 17 | -1.38 | 0.339 | 0.22 | -1.56 | 0.310 | 0.25 |
| | 16 - 17 | 1.71 | 0.218 | -0.27 | 0.35 | 0.824 | -0.05 |

Table S5: Bootstrap data corresponding to figure 4: PLSC results from for positive psychotic symptoms. The table shows bootstrap mean and 5 to 95 percentiles of behavior weights and brain weights.

| Weights type | item | mean | 5th percentile | 95th percentile |
|---|---|---|---|---|
| Behavior: SIPS Positive Symptoms | P1:Del (22q) | 0.64 | 0.403 | 0.89 |
| | P2:Susp (22q) | 0.51 | 0.307 | 0.72 |
| | P3:Gand (22q) | 0.32 | 0.115 | 0.53 |
| | P4:Hall (22q) | 0.49 | 0.238 | 0.76 |
| | P5:DisCom (22q) | 0.62 | 0.416 | 0.83 |
| iCAPs activation duration | dACC/dlPFC (5) | 0.38 | -0.002 | 0.78 |
| | PrimVis2 (8) | 0.03 | -0.246 | 0.33 |
| | FPN (9) | 0.59 | 0.144 | 1.05 |
| | aDMN (10) | -0.15 | -0.490 | 0.21 |
| | pDMN (11) | -0.01 | -0.356 | 0.33 |
| | SM (6) | 0.10 | -0.288 | 0.49 |
| | iTEMP/FUS (14) | 0.66 | 0.347 | 0.97 |
| | AMY/HIP (15) | 0.14 | -0.271 | 0.57 |
| | OFC (16) | 0.09 | -0.301 | 0.50 |

| Weights type | item | mean | 5th percentile | 95th percentile |
|---|---|---|---|---|
| Behavior: SIPS Positive Symptoms | P1:Del (22q) | 0.69 | 0.468 | 0.90 |
| | P2:Susp (22q) | 0.54 | 0.318 | 0.76 |
| | P3:Gand (22q) | 0.18 | -0.031 | 0.40 |
| | P4:Hall (22q) | 0.44 | 0.211 | 0.67 |
| | P5:DisCom (22q) | 0.54 | 0.324 | 0.77 |
| anti-coupling duration with dACC/dlPFC (5) | AUD (3) | 0.03 | -0.320 | 0.38 |
| | FPN (9) | 0.41 | 0.005 | 0.86 |
| | AUD/SM (13) | 0.31 | -0.193 | 0.83 |
| | iTEMP/FUS (14) | 0.88 | 0.575 | 1.22 |
| | AMY/HIP (15) | 0.01 | -0.440 | 0.50 |

Table S6: Bootstrap data corresponding to figure 5: PLSC results from for anxiety. The table shows bootstrap mean and 5 to 95 percentiles of behavior weights and brain weights.

| Weights type | item | mean | 5th percentile | 95th percentile |
|---|---|---|---|---|
| Behavior: | HC | 0.74 | 0.406 | 1.07 |
| CBCL/ABCL Anxiety | 22q11DS | 0.87 | 0.545 | 1.23 |
| iCAPs activation | dACC/dlPFC (5) | 0.05 | -0.301 | 0.42 |
| duration | PrimVis2 (8) | 0.27 | -0.085 | 0.63 |
| | FPN (9) | -0.12 | -0.494 | 0.28 |
| | aDMN (10) | -0.39 | -0.770 | -0.02 |
| | pDMN (11) | -0.16 | -0.536 | 0.20 |
| | SM (6) | -0.18 | -0.510 | 0.14 |
| | iTEMP/FUS (14) | 0.68 | 0.304 | 1.03 |
| | AMY/HIP (15) | 0.41 | 0.033 | 0.78 |
| | OFC (16) | -0.20 | -0.479 | 0.08 |
| Weights type | item | mean | 5th percentile | 95th percentile |
| Behavior: | HC | -0.11 | -0.350 | 0.17 |
| CBCL/ABCL Anxiety | 22q11DS | 1.05 | 0.688 | 1.44 |
| Positive coupling | LAN (4) | 0.35 | 0.014 | 0.67 |
| duration with | dACC/dlPFC (5) | 0.75 | 0.400 | 1.05 |
| AMY/HIP (15) | PREC/vDMN (7) | -0.06 | -0.375 | 0.25 |
| | FPN (9) | 0.29 | -0.024 | 0.60 |
| | aDMN (10) | -0.32 | -0.592 | -0.04 |
| | iTEMP/FUS (14) | 0.31 | -0.071 | 0.67 |

Table S7: Test statistics for group comparison of iCAPs' activation duration in high motion vs. low motion healthy subjects (shown in Supplementary Figure S5. P-values are corrected for multiple comparisons using false discovery rate; age and gender were included as covariates.

| iCAP | t-statistic | p-value | effect size (Cohen's d) |
|---|---|---|---|
| PrimVIS1 (1) | -0.27 | 0.795 | 0.06 |
| SecVIS (2) | 1.89 | 0.211 | -0.41 |
| aIN (3) | -2.01 | 0.205 | 0.44 |
| LAN (4) | 0.71 | 0.669 | -0.15 |
| dACC/dlPFC (5) | -0.60 | 0.669 | 0.13 |
| SM (6) | -1.57 | 0.294 | 0.34 |
| PREC/vDMN (7) | -0.63 | 0.669 | 0.14 |
| PrimVis2 (8) | 0.36 | 0.795 | -0.08 |
| FPN (9) | 1.39 | 0.330 | -0.30 |
| aDMN (10) | 2.70 | 0.071 | -0.59 |
| pDMN (11) | 2.71 | 0.071 | -0.59 |
| VSN (12) | 0.75 | 0.669 | -0.16 |
| AUD/SM (13) | -1.25 | 0.367 | 0.27 |
| iTEMP/FUS (14) | -1.37 | 0.330 | 0.30 |
| AMY/HIP (15) | -2.08 | 0.205 | 0.45 |
| OFC (16) | -1.81 | 0.211 | 0.39 |
| PFC (17) | -0.26 | 0.795 | 0.06 |
